# A better brain? Alternative spliced STIM2 in hominoids arises with synapse formation and creates a gain-of-function variant

**DOI:** 10.1101/2023.01.27.525873

**Authors:** Vanessa Poth, Hoang Thu Trang Do, Kathrin Förderer, Thomas Tschernig, Dalia Alansary, Volkhard Helms, Barbara A. Niemeyer

## Abstract

Balanced Ca^2+^ homeostasis is essential for cellular functions. STIM2 mediated Store-Operated Ca^2+^ Entry (SOCE) regulates cytosolic and ER Ca^2+^ concentrations, stabilizes dendritic spine formation and drives presynaptic spontaneous transmission and ER stress in neurons. Recently identified alternative spliced variants expand the STIM protein repertoire, uncover unique functions and facilitate our understanding of tissue specific regulation of SOCE. Here, we describe an addition to this repertoire, a unique short STIM2 variant (STIM2.3/STIM2G) present only in old world monkeys and humans with expression in humans starting with the beginning of brainwave activity and upon synapse formation within the cerebral cortex. In contrast to the short STIM1B variant, STIM2.3/STIM2G increases SOCE upon stimulation independently of specific spliced in residues. Basal cluster formation is reduced and analyses of several additional deletion and point mutations delineate the role of functional motifs for Ca^2+^ entry, NFAT activation and changes in neuronal gene expression. In addition, STIM2.3/STIM2G shows reduced binding and activation of the energy sensor AMPK. In the context of reduced STIM2.3 splicing seen in postmortem brains of patients with Huntington’s disease, our data suggests that STIM2.3/STIM2G is an important regulator of neuronal Ca^2+^ homeostasis, potentially involved in synapse formation/maintenance and evolutionary expansion of brain complexity.

## Introduction

Alternative splicing takes place in about 95% of human genes [1, 2]. With the rapid development of RNA microarrays, bulk RNA sequencing and sc-RNA sequencing technologies, the complexity of regulated and dynamic alternative splicing is just beginning to be understood [3-5] and it becomes increasingly evident that resulting alternate protein functions can shape both the physiology and pathophysiology of organisms [6]. Proteins derived from alternative splicing can alter infrared sensing in bats [7], sex determination in fruit flies [8] as well as change cell proliferation, cancer and neurological disorders [6]. So perhaps it is not surprising that a ubiquitous mechanism essential for regulation of cellular Ca^2+^ homeostasis and transcription factor activation is subject to cell-type specific alternative splicing. Store-operated Calcium Entry (SOCE) is triggered when activation of cell surface receptors induce depletion of Ca^2+^ from the endoplasmic reticulum (ER). The decrease in luminal Ca^2+^ is differentially sensed by STIM proteins with STIM2 -having a reduced EF-hand affinity-responding to small changes in intraluminal Ca^2+^ whereas STIM1 requires more substantial depletion [9, 10]. Activated STIM molecules gather at ER-PM junctions where they bind and activate ORAI ion channels residing in the plasma membrane (reviewed in [11]). STIM genes (STIM1 and STIM2), have a related genomic structure with 12 conventional exons [12] and long introns harboring partly unknown small exons (reviewed in [13]), thus may utilize alternate splicing to adapt SOCE to cell type specific needs. The first described STIM1 splice variant shows an alternatively spliced extension of exon 11, leading to the longer protein variant STIM1L found in skeletal muscle [14]. A dramatic switch in STIM2 function can be seen with alternative exon inclusion into the region encoding the ORAI ion channel activating domain (SOAR/CAD), with splice inclusion reverting STIM2 from a channel activator to an inhibitor of channel function [15, 16]. Besides the inhibitory splice variant STIM2.1 (STIM2ß), we have recently described how two alternate tissue specific splice variants of STIM1, which although modifying SOCE to a much lesser degree, can profoundly influence the efficacy of synaptic transmission in a frequency dependent manner, in the case of the neuronal-specific STIM1B [17], or differentially affect gating of ORAI1, protein interactions and NFAT translocation in the case of the more broadly expressed STIM1A, [18]. This same variant (STIM1A) has also been described as STIM1ß and shown to alter glioblastoma proliferation and wound healing [19], although both of these reports postulate different molecular mechanisms leading to altered cellular function.

With the discovery of a neuron-specific STIM1 splice variant [17] and the finding that STIM2 is prominently expressed in the brain, where it contributes to hypoxia induced neuronal death [20], but also stabilizes dendritic spine formation and protects spines from amyloid synaptotoxicity [21, 22] as well as positively affects spontaneous excitatory neurotransmission and drives Synaptotagmin7 dependent neurotransmitter release [23], the aim of this study was to investigate expression and function of the alternate STIM2 splice variant STIM2.3/STIM2G.

## Results

During initial screening for novel STIM2 splice variants [15], we detected an alternative spliced exon within STIM2’s critical channel activating region, leading to slightly longer STIM2.1 (NM_001169118.2) and expressed in several cell types. However, we were unable to detect any significant amount of the predicted variant STIM2.3 (STIM2G) (NM_001169117.2) in lymphocytes or cell lines. Detailed analysis of novel splice variants of STIM1, meanwhile, revealed a new STIM1 variant with an alternative exon inserted between conventional exons 11 and 12, resulting in the short protein variant STIM1B, present in neuronal cells [17]. Although the alternate STIM1 exon is not conserved in STIM2, we find that the putative alternate exon 13 also resides within the intronic region between conventional exons 11 (now 12) and 12 (now 14), located on chromosome 4p15.2; with variant STIM2.3 described according to HGVS as: NC_000004.12(NM_020860.4): c.1763_1764 ins1764-1518_1764-1451, resulting in an mRNA coding sequence of 2061 nt, compared to 2502 nt for Stim2.2 (Fig. 1A). The inserted exon results in termination of the protein after 686 amino acids and contains 12 unique amino acids. The abbreviated protein lacks 159 of the original C-terminal residues including the serine/proline rich region (SP) as well as the C-terminal poly-basic domain (PBD) (Fig. 1B). In contrast to all previously analyzed variants (STIM2.1/STIM2ß, STIM1B, STIM1A [18], this splice event evolved more recently and the inserted unique protein domain can only be found in *Catarrhini* (old world monkeys): *Theropithecus gelada* (gelada baboons) and *Hominoids (*apes), implying a selective advantage that arose ∼20 Million years ago (Fig. 1C). In contrast to the alternate exon B for STIM1, exon 13 also contains a short poly-adenylation site. Using conventional and splice-specific primers (Fig. 2A) as well as counting reads derived from RNA seq data (exemplary reads shown in Fig. 2B), we detected STIM2.3 (STIM2G) in postmortem human brain probes extracted from different brain regions (Fig. 2C), with variability as expected from tissues derived from different postmortem times but potentially with a slightly higher expression of both *STIM2* and *STIM2*.*3 (STIM2G)* in female compared to male donors by qRTPCR analysis (Fig. 2C). We also used commercially available qRTPCR primers for general *STIM1* and *STIM2* detection and observed slightly increased *STIM1* versus *STIM2* expression in cerebellum. In addition, we also find *ORAI2* as the most abundant *ORAI* isoform in human cerebellum (Fig. 2D). We next quantified the fraction of exon 13 inclusion, showing a slightly increased abundance in female donors (Fig. 2E). However, exon read analysis using published RNA seq data of postmortem tissues from different sources, while confirming inclusion of exon 13 in ∼20% of all reads, did not reveal gender differences (Fig. 2F). To estimate expression during early development, we analyzed RNA seq data obtained from the human developmental biology resource (HDBR) database to determine exon inclusion at ∼7 (Carnegie stage 22) and 9 weeks post conception and find significantly reduced exon inclusion during fetal brain development, indicating a potential role of STIM2.3 in synapse formation/pruning at later developmental stages (Fig. 2G). We did not detect splice inclusion in samples (0/4) derived from astrocytes differentiated from iPSCs, indicating that STIM2.3 splicing may indeed be neuron-specific.

**Figure 1.**
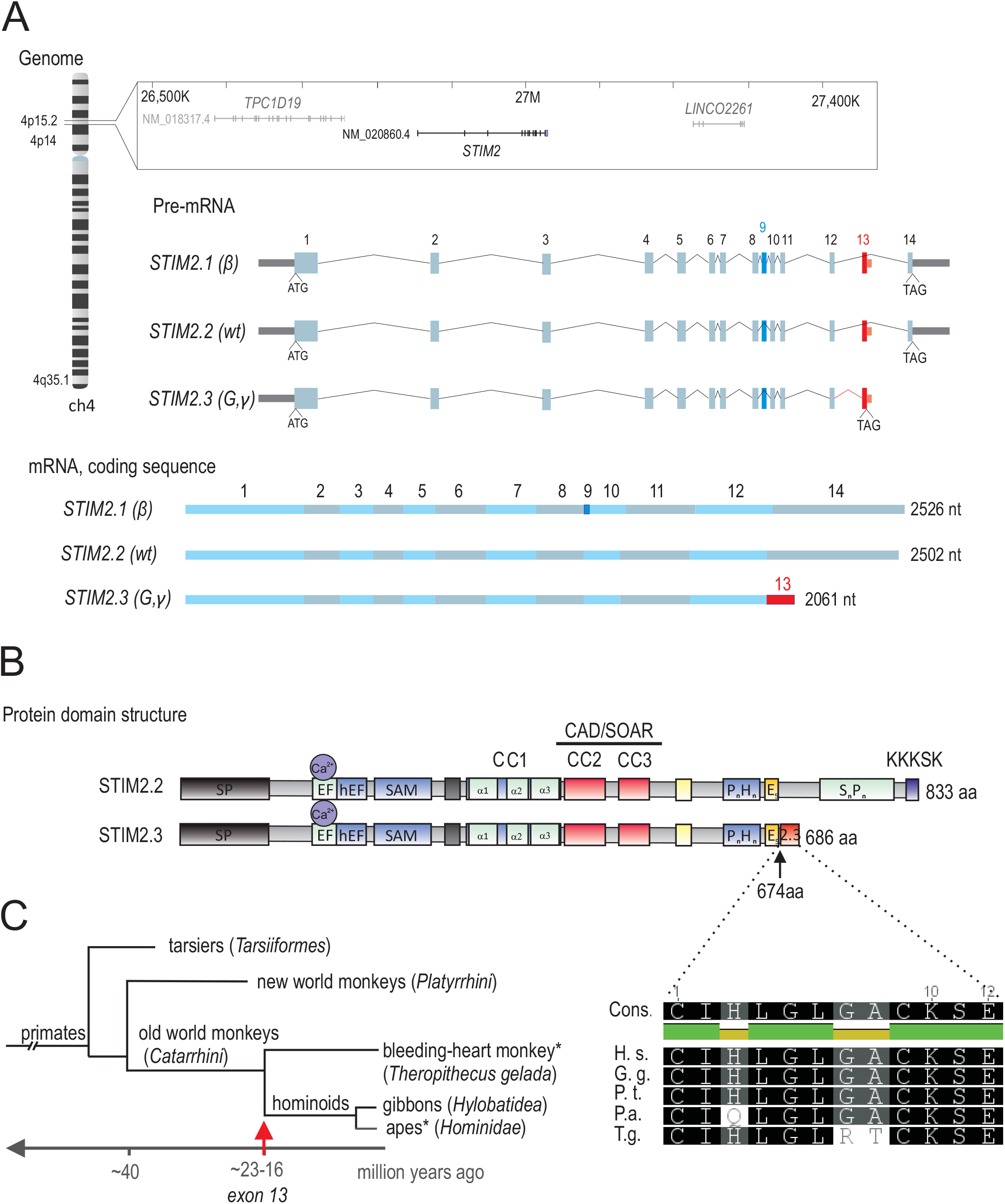
A. Genomic context and gene structure of human *STIM2*. Predicted pre-mRNA of the isoforms *STIM2*.*1* (ß), *STIM2*.*2* (wt) and *STIM2*.*3* (C). Translation start (ATG) and stop (TAG) codons are indicated. Conventional exons are depicted in light blue, STIM2.1 specific exon (9) is highlighted in dark blue and STIM2.3 specific exon (13) is highlighted in red. Lines indicate intronic regions. Coding sequence of *STIM2*.*1, STIM2*.*2* and *STIM2*.*3*. Alternating light blue and grey rectangles indicate individual exons. STIM2.3-specific exon (13) is highlighted in red. B. Schematic protein structure of STIM2 displaying functional domains. STIM2.3-specific domain (red) is inserted after poly-E domain (E_5,_ yellow) at aa 674. C. Phylogenetic tree. Red arrow indicates evolutionary splice event of exon STIM2.3. Evolutionary conservation of domain STIM2.3 in Homo sapiens (H. s.), Gorilla gorilla gorilla (G. g.). Pan troglodytes (P. t.). Pongo abelii (P. a.) and Theropithecus gelada (T. g.). Identical residues are within black boxes.

**Figure 2.**
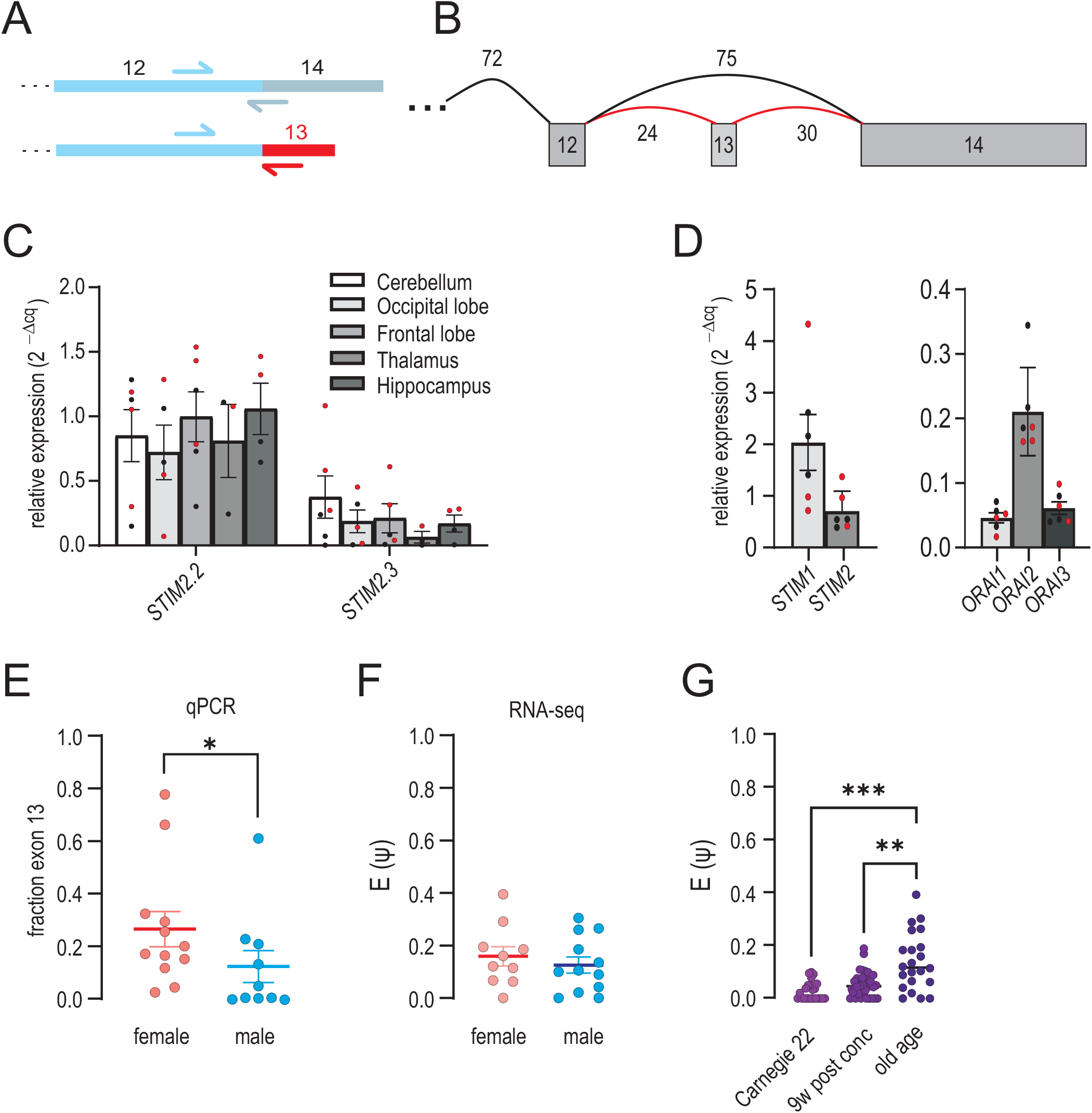
A. Schematic representation of primer annealing sites on coding mRNA. B. Representative RNA Seq read numbers of splicing of exon 13. C. Relative expression (2^-(ΔCq)^ ± SEM) of *STIM2*.*2* and *STIM2*.*3* using splice specific primers in different postmortem human brain regions as indicated derived from six different donors. Male donors are shown in black, female donors are shown in red. D. Relative expression (2^-(ΔCq)^ ± SEM) of both *STIM1* and *STIM2* and *Orai1-3* in cDNA of postmortem human cerebellum derived from six different donors. Male donors are shown in black, female donors are shown in red. E. Quantification of fraction exon 13 normalized to sum of splice-specific and wt-specific STIM2 measured in D in female or male donors. Quantification of spliced-in exon 13 in female or male donors using RNA Seq data of cerebellum and frontal cortex. F. Quantification of spliced-in exon 13 in different developmental stages of female and male donors using RNA Seq data of cerebellum, cortex and telencephalon. G.

### STIM2.3 (STIM2G) is a gain-of-function variant

To assess functional differences of the alternative STIM2 splice variants, we expressed YFP tagged isoforms in cells lacking endogenous levels of STIM1 or STIM2 [24]. While a vector only transfected control displayed low resting Ca^2+^ levels and no store-operated re-entry of Ca^2+^ after thapsigargin (Tg) induced store depletion, expression of wildtype STIM2 (STIM2.2) in the absence of STIM1 only slightly raised basal Ca^2+^ levels and recovered a small re-entry after re-addition of Ca^2+^ to the medium (Fig. 3A). Surprisingly, the C-terminally-deleted variant STIM2.3 (STIM2G), while also showing a higher basal Ca^2+^ compared to controls, but similar to STIM2.2, showed increased rates, peak and plateau of SOCE (Fig. 3A,B). To validate these results in a cell line more closely resembling neurons and with endogenous levels of STIM1 present, we generated SH-SY5Y cells lacking only STIM2 by Crispr/Cas9 (Fig. S1A). Compared to HEK293 cells, SH-SY5Y cells show higher expression of endogenous ORAI2 (Fig. S1B). Re-expression of either STIM2 variant in these cells and calibration of the Fura2 ratios to yield absolute [Ca^2+^] values uncovered a small increased basal Ca^2+^ concentration upon STIM2.3 (STIM2G) expression when compared to STIM2.2 and otherwise confirmed the phenotype of increased SOCE seen in the double knock-out HEK293 background (Fig. 3C,D). As expected with endogenous levels of STIM1 present, SOCE in the absence of STIM2 was not eliminated. Equal protein expression was controlled by recording the tagged YFP-fluorescence of measured cells (Fig. 3E) and in addition by determining protein expression of constructs in which the N-terminal YFP was replaced by an HA tag, yielding an expected molecular mass of ∼ 68 kDa for STIM2.3 (Fig. 3F). As it is unlikely that only one species of a STIM2 variant exists in a given cell, we checked if STIM2.3 increases SOCE also in the presence of STIM2.2. Co-expression of YFP-tagged STIM2.2 or STIM2.3 with mkate2-tagged STIM2.2 or STIM2.3 demonstrates that the presence of STIM2.3 has a reduced but significant amplifying effect on rate, peak and plateau of SOCE (Fig. S1 C-D). Interaction analysis utilizing Bimolecular Fluorescent Complementation assays (BIFC) confirmed that STIM2.3 (STIM2G) forms heteromultimers with either STIM1 or STIM2 and also shows an increased interaction with ORAI2 when compared with STIM2.2 (Fig. S1E).

**Figure 3.**
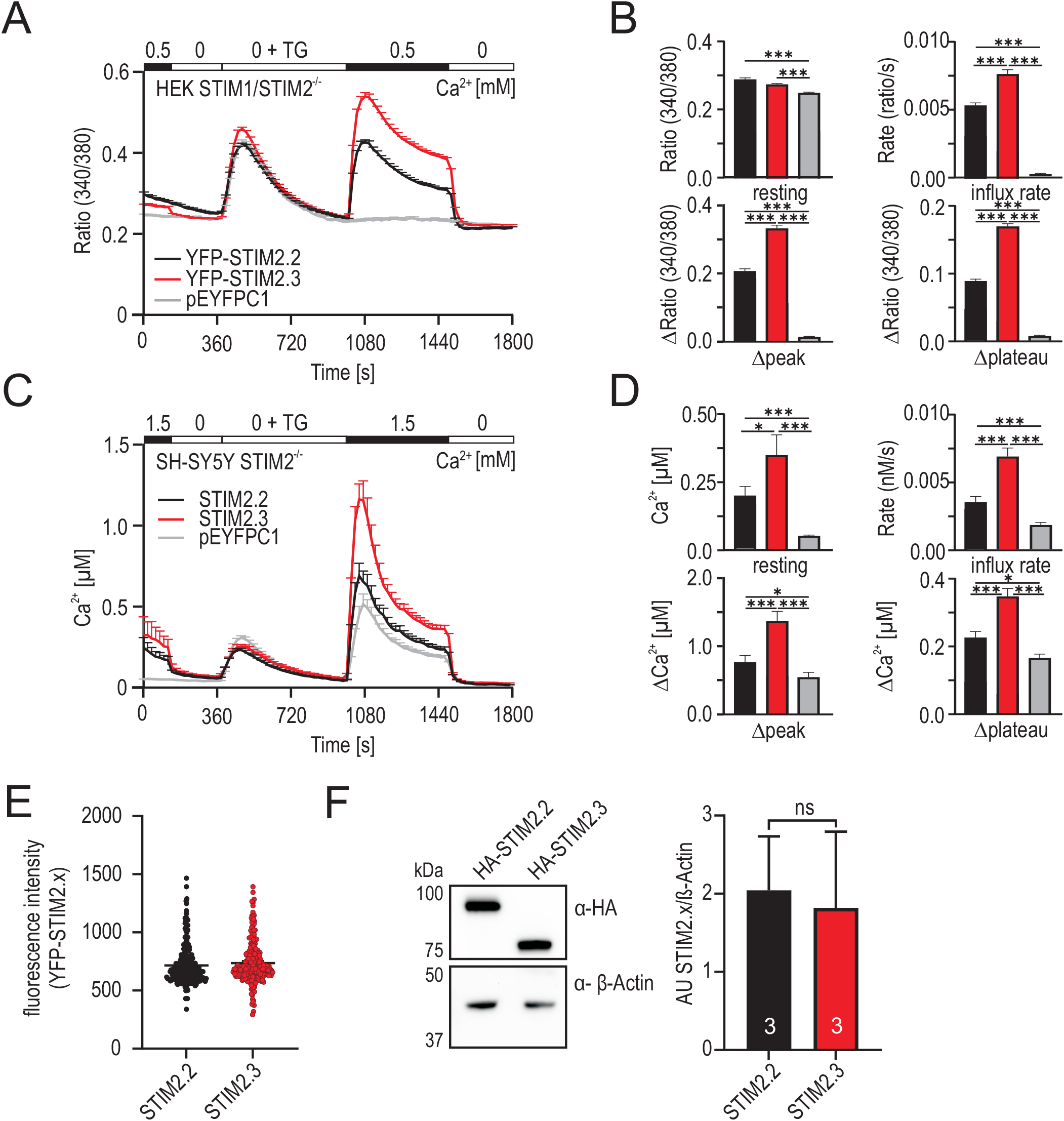
A. Traces showing average changes (mean + SEM) in intracellular Ca^2+^ (Ratio 340/380) over time in response to perfusion of different external [Ca^2+^]mM as indicated in the upper bar in HEK STIM1/2^-/-^ cells transfected with STIM2.2 (black, n=227), STIM2.3 (red, n=232) or vector only (grey, n=27). B. Quantification of changes in ratio of resting, influx rate (ratio/s), Δpeak and Δplateau measured in A. *** p < 0.001, Kruskal-Wallis ANOVA. C. Calibrated traces showing average changes (mean + SEM) in intracellular Ca^2+^ [μM] over time in response to perfusion of different external [Ca^2+^]mM as indicated in the upper bar after transfection with STIM2.2 (black, n=85), STIM2.3 (red, n=89) or vector only (grey, n=94) in SH-SY5Y STIM2^-/-^ cells. D. Quantification of changes in Ca^2+^ [μM] of resting, influx rate (nM/s), Δpeak and Δplateau measured in C. *** p < 0.001,* p < 0.05, Kruskal-Wallis ANOVA. E. Quantification of YFP fluorescence intensity from cells measured in A. F. Western blot showing HA-STIM2.2 and HA-STIM2.3 following heterologous expression in HEK STIM1/2^-/-^ cells (left); quantification from 3 independent transfections (mean ± SD).

### GOF phenotype does not require splice-specific residues and is not phenocopied by deletion of both EB binding sites and the PBD

To address whether the gain-of-function phenotype observed for STIM2.3 (STIM2G) is related to its spliced-in specific residues, we created an additional deletion mutant, which terminates the wild-type protein after residue 674, the last common residue. In addition, we also recreated a deletion of only the last five C-terminal lysine residues (D5K), which has previously been shown to result in a strong loss-of-function phenotype [25] (Fig. 4A). Indeed, when comparing SOCE signatures of these constructs in HEK DKO cells, we find that deletion of only the polybasic domain (D5K) reduces basal Ca^2+^ levels and SOCE to a significant extent, while the deletion at aa 674 phenocopies STIM2.3 (STIM2G), suggesting that deletion of a C-terminal inhibitory domain and not inclusion of splice-specific residues lead to STIM2.3 (STIM2G)’s gain-of-function phenotype (Fig. 4BC). To investigate whether retention of STIM2Δ5K to the microtubular network via its potential dual EB binding motifs (^773^SGIP^776^) and (^805^SSIP^808^) (numbering referring to long signal peptide (AAI36450.1) in contrast to the single site investigated by Pchitskaya et al. [26], is the cause of its strong loss-of-function phenotype and to test whether the lack of these motifs in STIM2.3 (STIM2G) is sufficient to explain its gain-of-function phenotype, we deleted both motifs in the background of STIM2.2 and STIM2.2Δ5K (Fig. 4A). While deletion of the EB binding motifs in the Δ5K mutant indeed rescued its SOCE phenotype to wildtype levels, STIM2.3 (STIM2G) still showed a significantly increased SOCE (Fig. 4DE). Interestingly, an additional deletion mutant (STIM2Δ711) did not phenocopy STIM2.3 (STIM2G) or STIM2Δ674, but showed wildtype-equivalent SOCE levels (Fig. S2), indicating a potential negative regulatory site between aa 674 to 711 within STIM2. However, deletion only of this region from full length STIM2 reduced and not increased SOCE (Fig. S2).

**Figure 4.**
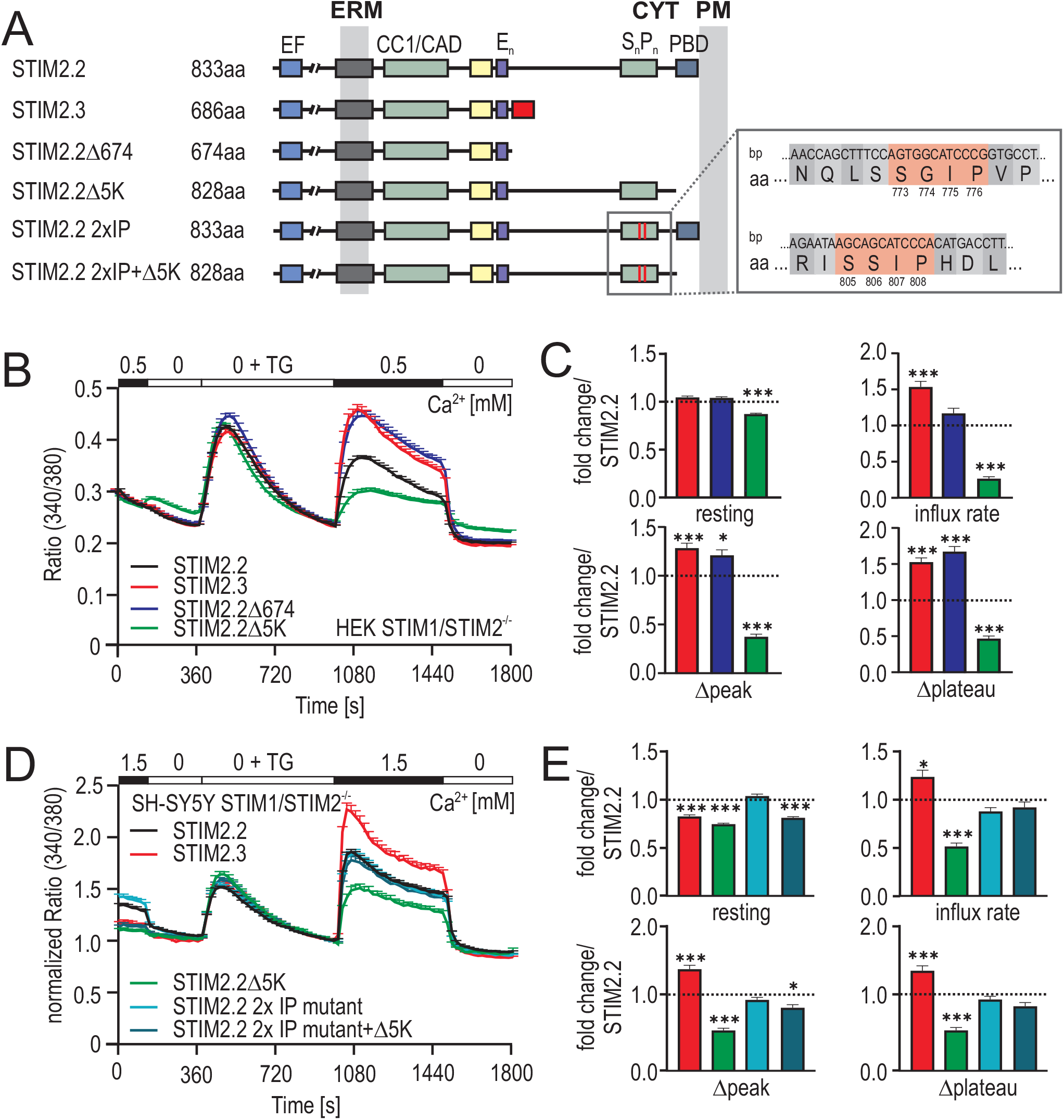
A. Schematic protein structure displaying functional domains of STIM2 splice variants and deletion mutants (Δ674/Δ5K/2xIP/2xIP+Δ5K). EB binding motifs are indicated in red within the S_n_/P_n_ rich region. Corresponding sequence (bp and aa) information is displayed in the grey box (right) with relevant aa indicated in red and corresponding aa position. B. Traces showing average changes (mean + SEM) in intracellular Ca^2+^ (Ratio 340/380) over time in response to perfusion of different external Ca^2+^ [mM] as indicated in the upper bar in HEK STIM1/2^-/-^ cells transfected with STIM2.2 (black, n=144), STIM2.3 (red, n=145), STIM2.2 Δ674 (blue, n=132) or STIM2.2 Δ 5K (green, n=164). C. Quantification of changes in ratio of resting, influx rate (ratio/s), Δpeak and Δplateau as fold change/STIM2.2 measured in B. *** p < 0.001; * p < 0.05, Kruskal-Wallis ANOVA. D. Normalized average traces showing changes (mean+SEM) in intracellular Ca^2+^ (Ratio 340/380) over time in response to perfusion of different external Ca^2+^ [mM] as indicated in the upper bar in SH-SY5Y STIM1/2^-/-^ cells transfected with YFP-STIM2.2 (black, n=198), YFP-STIM2.3 (red, n=130), YFP-STIM2.2Δ5K (green, n=214), YFP-STIM2.2 2xIP (light blue, n=158) or STIM2.2 2xIP+Δ5K (petrol blue, n=168). E. Quantification of changes in resting Ca^2+^, influx rate, Δpeak and Δplateau measured in B as fold change normalized to STIM2.2. *** p< 0.001, ** p< 0.01, * p< 0.05; Kruskal-Wallis ANOVA.

### Localization and cluster formation

Expression of wildtype STIM2 (STIM2.2) has been shown to lead to pre-clustered and pre-activated SOCE complexes [15, 27]. We used mCherry tagged STIM1 and co-expression of either YFP-tagged STIM2.2 or STIM2.3 (STIM2G) to assess the degree of colocalization and pre-clustering of the isoforms. In the absence of stimulation, STIM2.3 (STIM2G) showed good colocalization with either STIM1 or STIM2 in ER regions close to the nucleus, however, STIM2.3 (STIM2G) was significantly less pre-clustered with STIM2.2 in plasma membrane near regions (Fig. 5A,B). The reduced pre-clustering of STIM2.3 can be phenocopied by either deletion of the PBD domain alone or by deletion at aa674, mimicking the SOCE phenotype of STIM2.3, thus demonstrating that the PBD alone stabilizes formation of pre-clusters (Suppl Fig. S3A). Strong Tg-induced store depletion abolished differences in colocalization (Fig. 5C,D) and no differences concerning cluster sizes could be detected when comparing STIM2.3 (STIM2G) co-clusters with either STIM1 or STIM2.2 (Fig. 5E).

**Figure 5.**
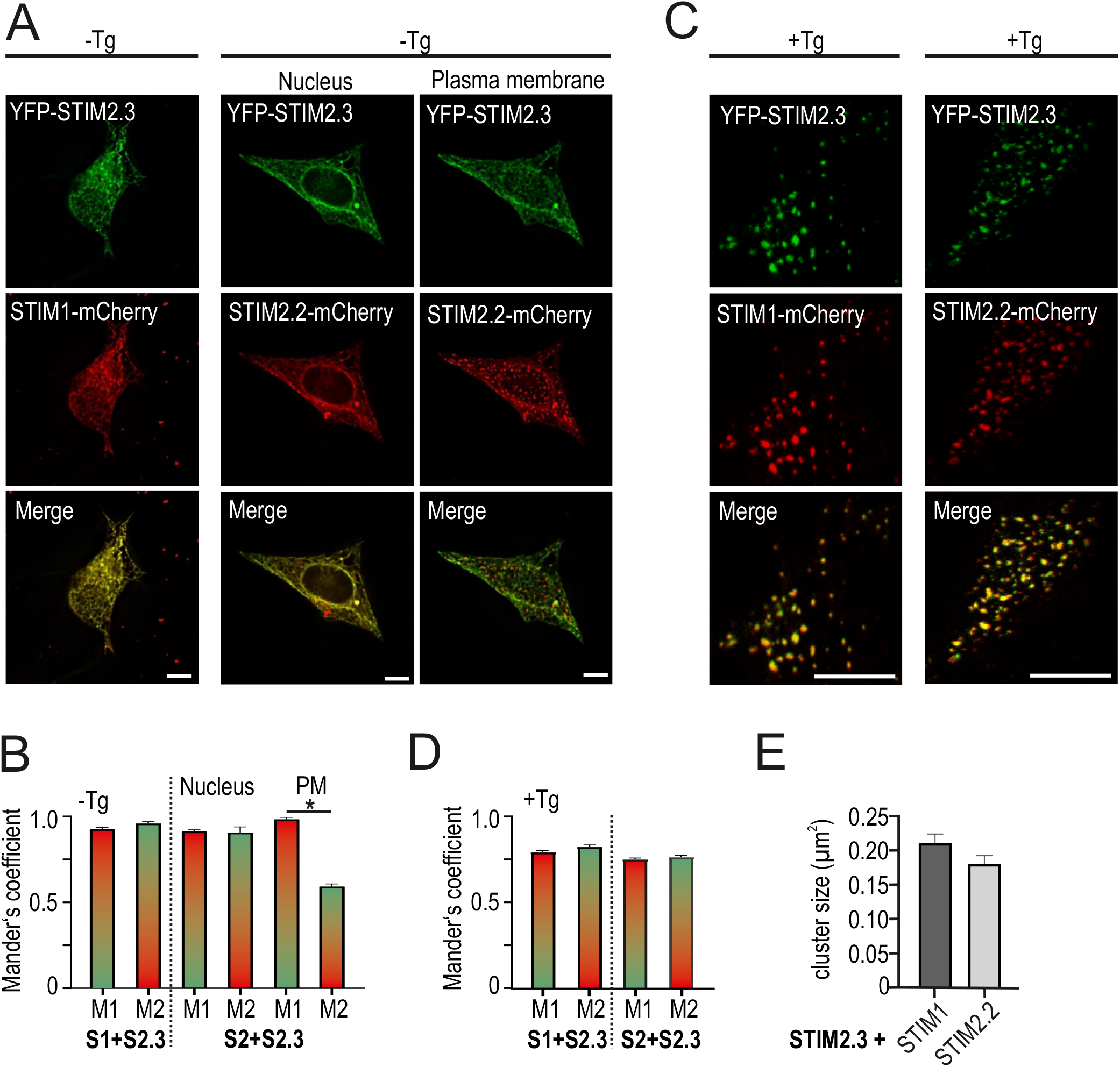
A,C. Representative images of HEK STIM1/2^-/-^ cells co-transfected with YFP-STIM2.3 (green) and STIM1-mCherry (red) (left) or STIM2.2-mCherry (red) (right) and merged images before (A) and after (C) stimulation with thapsigargin. Scale bar indicates 10μM. B,D. Co-localization analysis of cells from (A,C). For each condition 5-45 cells from 3 independent transfections were analyzed using Mander’s overlapping coeffients (M1, M2) (mean + SEM): S1+S2.3 –Tg M1:0.91 M2: 0.95; S1+S2.3 +Tg M1: 0.78 M2: 0.81; S2.2+S2.3 –Tg nucleus M1: 0.90 M2: 0.89; S2.2+S2.3 –Tg plasma membrane M1: 0.97 M2: 0.58; S2.2 +S2.3 +Tg M1: 0.74 M2: 0.75. E. Quantification of cluster size (mean ± SEM) from cells measured in (C). For each condition 16-45 cells from 3 independent transfections were analyzed.

We next asked how and if the degree of Tg induced cluster formation and cluster size depends on the presence or absence of either the PBD, the EB binding sites or on the larger deletion found in STIM2.3 (STIM2G). Cells transfected with YFP tagged constructs and PH-PLC-mCherry to indicate the plane of the PM for TIRF microscopy and to measure regions of high PIP_2_ were imaged before and after Tg (exemplary images shown in Figure 6A). Despite the ability of STIM2.3 (STIM2G) to increase SOCE, STIM2.3 (STIM2G), but also STIM2Δ5K as well as STIM2 2xIP+Δ5K all showed a significant reduction in mean cluster sizes and cluster intensities (Fig 6B,C), demonstrating that cluster size is not a valid predictor of SOCE size (Fig. 6AB). In addition, as recently shown [28], PI4P and not PIP_2_ critically regulates STIM protein attachment to the plasma membrane.

**Figure 6.**
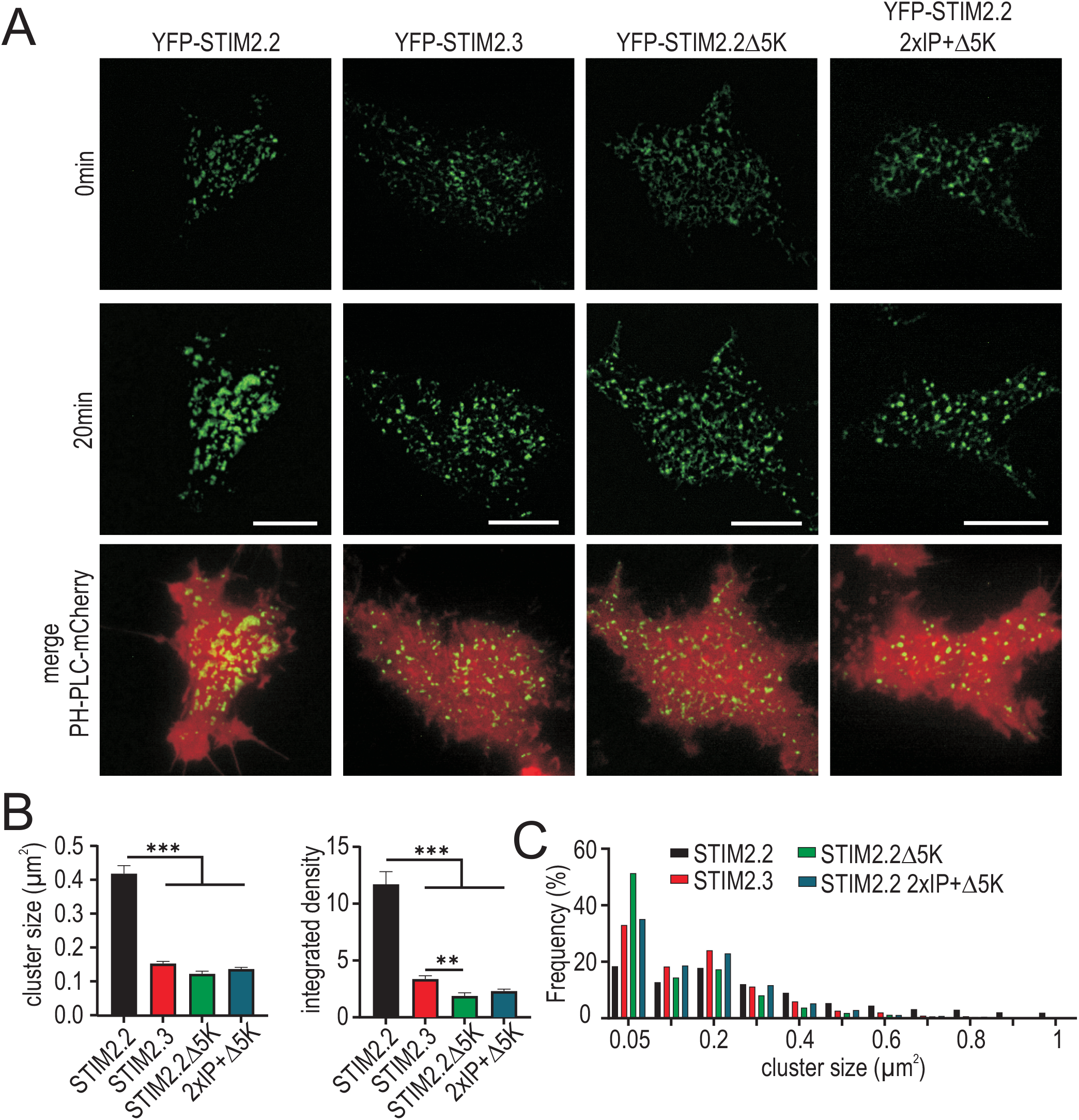
A. Representative images of HEK STIM1/2^-/-^ cells co-transfected with PH-PLC-mCherry (red, merge) and YFP-STIM2.2, YFP-STIM2.3, YFP-STIM2.2Δ5K or YFP-STIM2.2 2xIP+Δ5K before (0 min) and after (20 min) stimulation with thapsigargin. Scale bar indicates 10μM. B. Quantification of cluster size and integrated density from cells measured in (A). For each condition 34-50 cells from 3 independent transfections were analyzed. ** p < 0.02, *** p < 0.001, Kruskal-Wallis ANOVA. C. Cluster size distribution (frequency %) from cells measured in (A).

### Splice isoform dependent effects on NFAT activation

To assess downstream effectors of SOCE, we investigated the ratio of cytosolic to nuclear GFP-tagged NFATc1. mKate2-tagged STIM2 constructs were co-transfected with GFP tagged NFAT and the nuclear to cytosolic ratios of NFAT were analyzed before and after stimulation. As expected from its higher degree of plasma membrane pre-clustered protein, expression of STIM2, but not of STIM2.3 (STIM2G) or deletion mutants without a PBD, showed a high degree of basal NFAT translocation (Fig. 7AB) in the absence of stimulation. While store depletion did not lead to a much larger translocation in the case of STIM2, STIM2.3 (STIM2G) and the 2xIP+Δ5K mutant of STIM2 showed significant changes in NFAT translocation ratios upon stimulation (Fig. 7B). Here, it also becomes apparent that the tug-of-war between PM attachment via the PBD and retention to the MT via the IP motifs, reflects the ability to initiate NFAT translocation, even in the absence of STIM1. While STIM2Δ5K is unable to cause NFAT translocation, additional mutations within the IP motifs is sufficient to restore SOCE (Fig. 4DE) as well as recover NFAT translocation (Fig. 7AB), confirming the submembrane microdomain as an essential docking site to initiate NFAT translocation [29]. To cleanly correlate global changes in [Ca^2+^]_i_ and NFAT translocation, we also recorded SOCE with the same time course and parameters used for the NFAT translocation assays and quantified resting Ca^2+^ and total stimulated Ca^2+^ entry as area under the curve (AUC) during stimulation. Fig. 7C shows mean global Calcium responses to a 30 min stimulation. While vector only and STIM2Δ5K transfection showed neither basal Ca^2+^ entry nor a sustained SOCE and did not enable NFAT translocation, the other constructs reached a critical sustained [Ca^2+^]_i_ threshold to initiate nuclear translocation. These results also demonstrate the non-linearity between SOCE and endpoint (30 min) NFAT translocation, as despite different degrees of Calcium entry (Fig. 7D), STIM2.2, STIM2.3 (STIM2G) and STIM2 2xIP+Δ5K all result in a similar degree of translocation 30 min after stimulation (Fig. 7B). The finding that STIM2.3 (STIM2G), or the STIM2.2 2xIP Δ5K mutant despite also raising basal [Ca^2+^]_i_ do not lead to pre-stimulus-translocated NFAT is congruent with their significantly decreased degree of ER-PM junction pre-clustering (Fig. 6) and implies that pre-cluster size correlates to an increased recruitment of the ORAI1-AKAP-NFAT signalosome.

**Figure 7.**
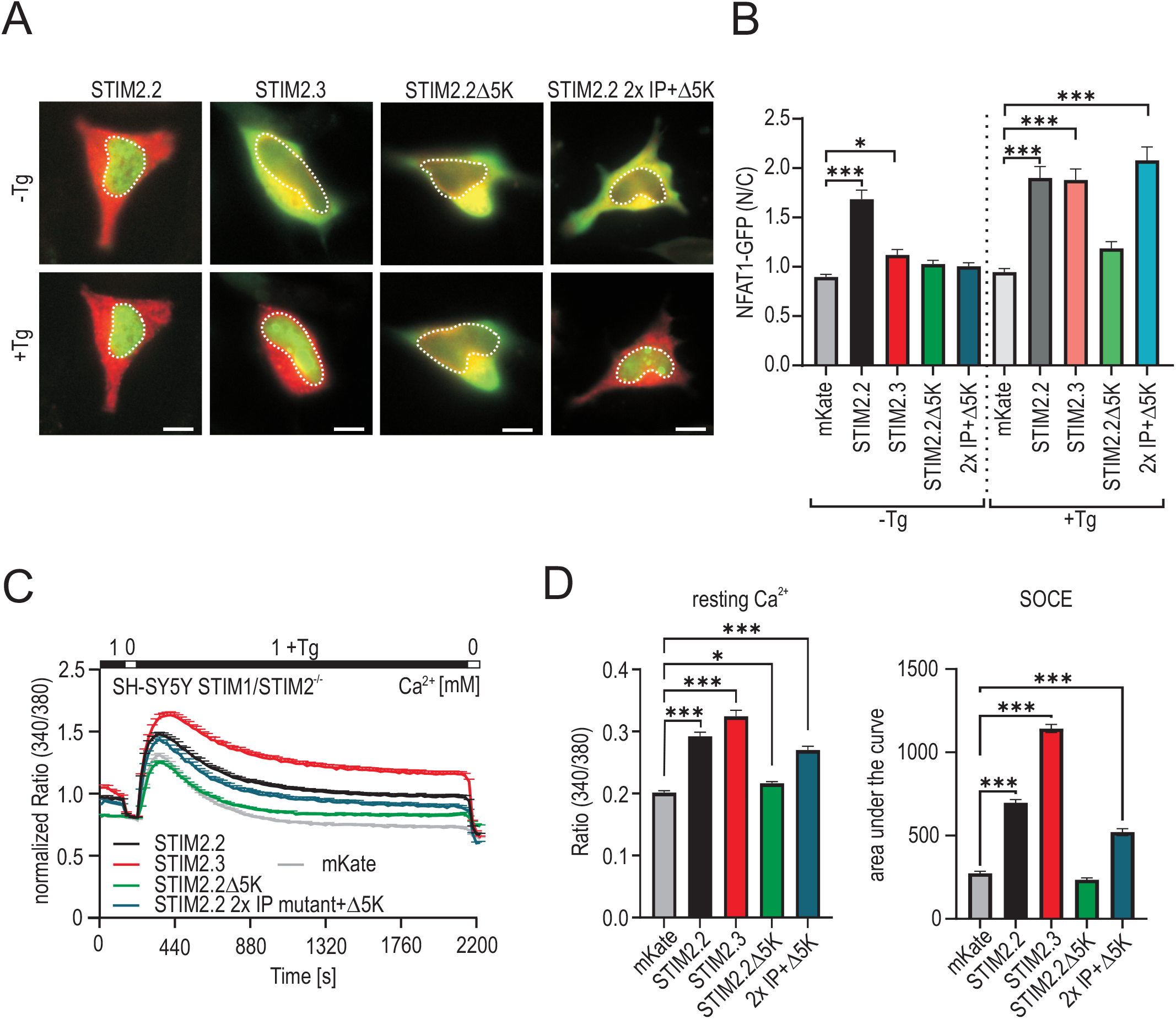
A. Representative images of SH-SY5Y STIM1/2^-/-^ cells co-transfected with NFAT1-GFP (green) and mKate2-STIM2.2 (red), mKate2-STIM2.3 (red), mKate2-STIM2.2Δ5K (red) or mKate2-STIM2.2 2xIP+Δ5K (red) before (-Tg) and after (+Tg) stimulation with thapsigargin (Tg). Scale bar indicates 10μM. B. Changes in ratio of nuclear/cytosolic (N/C) NFAT1-GFP before and after stimulation with thapsigargin in cells transfected as indicated. For each condition 39 -114 cells were analyzed from 3 independent transfections. *** p< 0.001, * p< 0.05; Kruskal-Wallis ANOVA. C. Normalized traces showing global changes in intracellular Ca^2+^ (Ratio 340/380) after perfusion of different external Ca^2+^ [mM] as indicated in the upper bar after transfection with mKate2-STIM2.2 (black, n=184), mKate2-STIM2.3 (red, n=160), STIM2.2Δ5K (green, n=156), STIM2.2 2xIP+Δ5K (dark blue, n=164) or vector only (grey, n=241) in SH-SY5Y STIM1/2^-/-^ cells. D. Quantification of resting Ca^2+^ (Ratio 340/380) and area under the curve from cells measured in (C). * p < 0.05, *** p < 0.001, Kruskal-Wallis ANOVA.

### STIM2.3 (STIM2G) reduces crosstalk and activation of the metabolism sensor AMPK

Besides its role in activating ORAI proteins, STIM2 can also function as a scaffolding protein, promoting assembly and activation of a trimeric complex between the energy sensor AMPK and CaMKKII [30]. Within STIM2, the C terminal region spanning aa 646-833 has been identified as the interacting domain with AMPK, thus, we investigated whether the truncated C terminus (last common aa 674) interferes with this interaction. Co-IP experiments in HEK STIM1/2 DKO cells transfected with HA-STIM2.2 or HA-STIM2.3 confirmed interaction of AMPK with STIM2.2 and revealed a reduced but not absent interaction of STIM2.3 with endogenous AMPKα, its catalytic subunit (Fig. 8AB). We also tested the ability of STIM2 to interact before and after store depletion and detected an increased interaction after Tg only in the case of STIM2.2 (Fig. 8C). Additionally, to monitor activation, we investigated the phosphorylation state of AMPKα at Thr172 within its activation loop, by a phospho-specific antibody. Concomitant to the reduced interaction we found reduced activation of AMPK (Fig. 8D), which is interesting as STIM2.3 increased SOCE. While endogenous AMPK can be detected in the lysates, we can only resolve active AMPK in the STIM2 bound fractions (compare middle panel of Fig. 8A,B). In neurons, hyperactive AMPK, indicating metabolic stress impairs neuronal polarization [31, 32]. To investigate in an unbiased manner whether expression of STIM2.3 might be beneficial for pro-neurogenic gene expression, we generated a STIM2.3-stably expressing SH-SY5Y cell line (E12.1) in a STIM2.2^-/-^ background. Expression of STIM2.3 was confirmed on protein level (Fig. S3B) and these cells show an increased SOCE (Fig. 8E) with significantly increased influx rate, Δpeak and Δplateau compared to SH-SY5Y STIM2^-/-^ cells (Fig. S3C). RNA Seq and subsequent Gene Ontology analysis of the STIM2.3-stable cell line compared to SH-SY5Y WT cells (expressing endogenous levels of STIM2.2) revealed a significant up-regulation of genes involved in neurogenesis, neuronal differentiation, axon development and guidance (Fig 8E and Fig. S3D).

**Figure 8.**
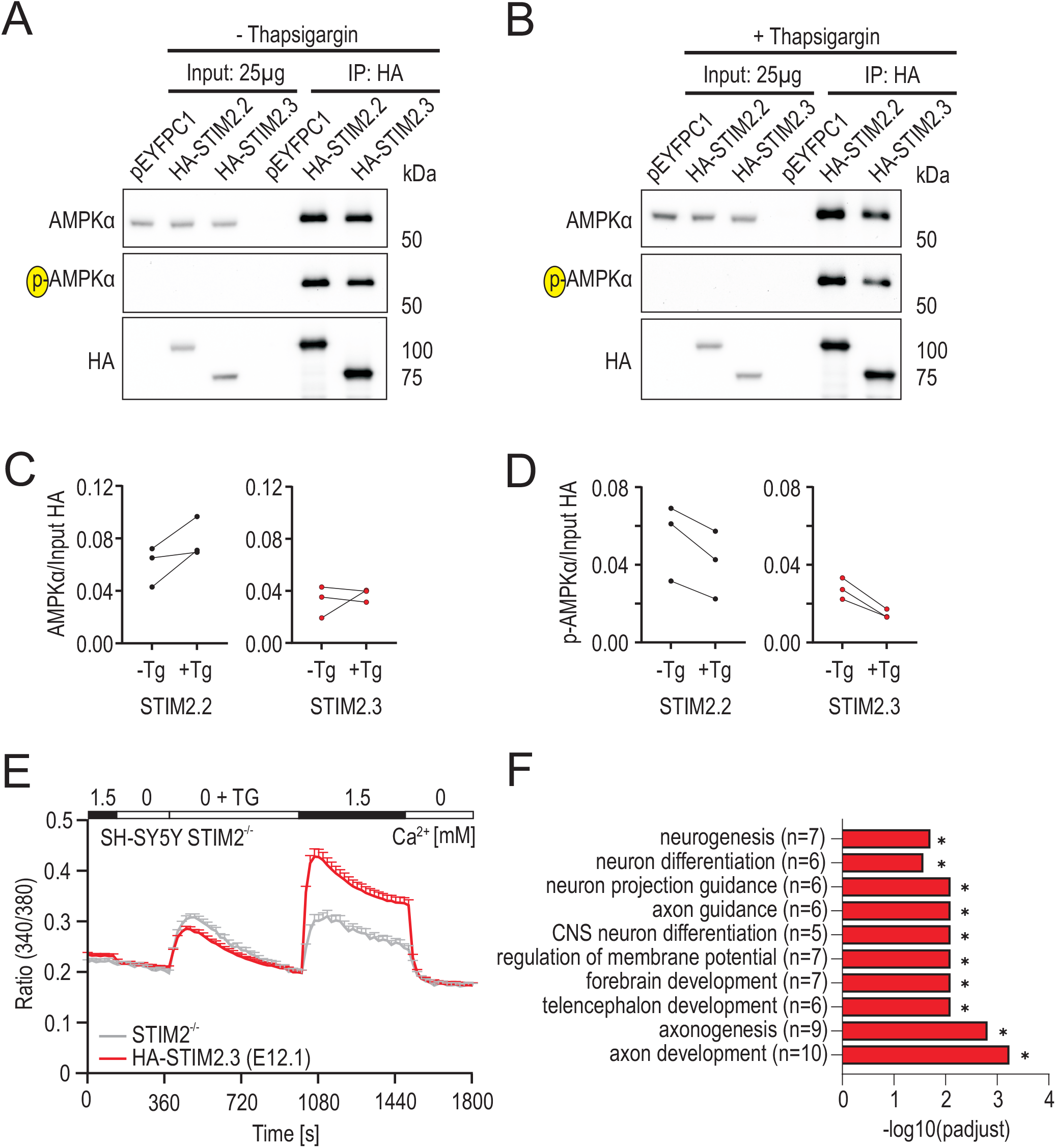
A, B. Co-immuniprecipitation of HA-STIM2.2 or Ha-STIM2.3 and endogenous AMPKα in HEK STIM1/2^-/-^ cells before (A) and after (B) stimulation with thapsigargin using anti-HA agarose. Membranes were incubated with the indicated antibodies. Cells expressing pEYFPC1 were used a control. C, D. Densitometric quantification of AMPK and p-AMPK signals before and after Tg treatment measured in (A,B) from 3 independent transfections. E. Average traces of changes in intracellular Ca^2+^ (Ratio 340/380) over time in response to perfusion with different external Ca^2+^ [mM] as indicated in the upper bar in SH-SY5Y STIM2^-/-^ (grey, n=31) and HA-STIM2.3 (E12.1, red, n=93) expressing clonal SH-SY5Y STIM2^-/-^ cell lines. F. Selected positive regulated Gene Ontology (GO) terms with the associated number of DEGs in the STIM2.3-stable cell line E12.1.

## Discussion

The present study identifies STIM2.3 (STIM2G) as a unique splice variant arising with the evolution of *Hominoids* and *Theropithecus*, indicating a potential selective advantage for the development or function of more complex brains. Indeed, a human-specific variant ARHGAP11B which evolved by a single splice site mutation has been shown be causative for neocortical expansion by enhancing basal progenitor generation [33-35]. Highest expression of the alternative spliced-in exon 13 is found within the cerebellum, however, we detected exon 13 expression in all investigated brain regions. Cerebellar expansion rate significantly increased relative to neocortex in the phylogenetic branch of apes compared to related non-ape branches, indicating an important role of cerebellar specialization in cognitive evolution including technical complexities such as production and use of tools and learning of complex motor skills [36], pointing to a potential contribution of STIM2.3. Generation of a stable STIM2.3 expressing cell line with subsequent analysis of its differentially expressed genes compared to the cell line expressing the non-spliced variant indeed indicates a significant upregulation of genes involved in neurogenesis, neuronal differentiation and axon development. STIM2.3 likely is of pathophysiological relevance as analysis of gene splicing in postmortem probes of Huntington’s disease patients, revealed that usage of the specific splice borders of alternative exon 13 showed a highly significant decrease in the PSI (percent splice inclusion: 0.175±0.035 wt vs 0.004±0.006 in HD) in striatum of HD patients compared to healthy controls [37], supplemental table S7). The percent striatal exon splice inclusion correlates well with data shown in Figure 2. A steady progression of motor dysfunction is a hallmark of Huntington’s disease, however, the molecular origins of the motor disfunction are not well understood [38] but correlate with a loss of cerebellar Purkinje neurons [39]. Utilizing patient-specific iPSC derived neurons, STIM2 has been found to mediate excessive SOCE in HD patient cells, potentially contributing to a juvenile form of HD, although splicing has not been investigated in this study [40].

We found that splice induced deletion of 159 residues of STIM2’s C-terminal domain did not reduce but instead increased SOCE. Deletion of only the PBD in full-length STIM2 significantly reduced SOCE confirming previous findings [25, 29]. In STIM1, deletion of only the PBD leads to retention of activated STIM1 to the microtubular network [41], suggesting a “tug of war” between PM attachment and MT retention for STIM1. Indeed, simultaneous mutation of both EB binding sites (2xIP) and the PBD (Δ5K) in STIM2 (2xIP+Δ5K) rescued the negative effect of STIM2 Δ5K on SOCE, demonstrating competitive functions of the PBD and EB binding motifs similar to previous findings for STIM1. However, STIM2.3 still showed increased SOCE compared to STIM2 (2xIP+Δ5K), while a deletion of residues 712 - 833 (STIM2.2Δ711) was similar to Wt or STIM2 (2xIP+Δ5K). These results suggest that the additional 37 amino acids compared to the mimicking mutant STIM2.2Δ674 may be involved in the STIM2.3-mediated SOCE phenotype. Surprisingly, internal deletion of these residues (Δ675-710), decreased SOCE. An increase in STIM1 function upon larger C-terminal deletion has recently been described for the TAM associated gain-of-function mutation STIM1^I484R^, leading to termination of translation at residue 502, abrogating MT binding sites and PBD [42]. Here, interpretation of its activated phenotype involves a potential disabled binding of the inhibitory domain (ID) of STIM1 to the CC1 region and disabled binding of STIM1 to SARAF, an accessory protein which maintains STIM proteins in the closed and inactive conformation [43, 44], or a combination of both. Interaction of SARAF with STIM1 entails a second binding site encompassing residues 490 to 530 of STIM1 [45], a domain that shows only 15% conserved residues between STIM1 and STIM2, but is not altered in STIM2.3 compared to STIM2, thus we believe that the mechanism leading to STIM2.3’s increased function is likely independent of SARAF. An additional difference between STIM1 and STIM2 lies in the different affinities of their respective intraluminal EF hands [10], resulting in considerable pre-clustering of STIM2 when expressed in HEKDKO cells. This pre-clustering is absent in STIM2.3 and also in all constructs lacking the PBD, confirming previous results with swapped C-terminal domains between STIM1 and STIM2 [46], indicating a dominance of STIM2’s PBD over its EF hand regarding pre-clustering. Additionally, we find that in the absence of endogenous STIM but with endogenous ORAI proteins present, all constructs lacking the PBD show reduced cluster sizes after store-depletion, confirming that the PBD enhances cluster formation and size/stability but also clearly indicates that cluster presence and size does not correlate well with ORAI activity.

Downstream effectors of SOCE include activation and nuclear translocation of the transcription factor NFAT, which also is of critical importance for neurons. Here, especially morphological remodeling of axon terminals during synaptogenesis requires CaN/NFAT-dependent gene expression [47]. In heterologous expression, STIM2.3 abrogated the permanently high nuclear NFAT translocation seen with STIM2.2 without store depletion, but translocated NFAT after store-depletion. In contrast to deletion of the PBD within STIM1, abrogating NFAT translocation also after stimulation [18, 25], we find for STIM2 that as long as intracellular Ca^2+^ reaches a critical threshold and ER-PM clusters are present, NFAT translocation is enabled, despite a lack of the PBD, in line with result showing that recruitment of AKAP requires the N-terminal anchoring domain within ORAI1 [48]. AMPK is a master regulator of cellular energy metabolism as it senses and monitors the concentration of the nucleosides AMP, ADP and ATP, which all bind competitively to the γ-subunit of the heterotrimeric αβγ-complex of the active enzyme. Previous work has shown that STIM2 interacts with AMPK [30, 49], and is also a substrate for AMPK regulation, leading to reduced SOCE activity by phosphorylation [50], reviewed in [51]. Our results show a reduced interaction and suggest a reduced activation of AMPK. Moreover, one of the AMPK substrates within STIM2, namely S680 is lacking in STIM2.3. Future work will delineate the role of STIM2 and STIM2.3’s regulation of AMPK activity in neurons and its role in synapse formation.

In summary, our work identifies STIM2 splicing as a promising regulator of neuronal SOCE. Its evolutionary late and brain specific addition to the STIM repertoire, with increased SOCE, reduced AMPK activation and differential activation of neurogenesis genes, leads us to hypothesize that STIM2.3 may be an important regulator of neuronal differentiation, dendritic spine formation or pruning and homeostatic synaptic activity, and that mis-splicing of STIM2 as observed in HD patients may contribute to neuronal degeneration.

## Methods

### Cell culture and transfection

Cell lines were cultured at 37°C, 5% CO_2_ in a humidified incubator in their respective medium (HEK STIM1/2^-/-^ Dulbecco’s modified Eagle’s medium, SH-SY5Y STIM2^-/-^ and STIM1/2^-/-^ Dulbecco’s modified Eagle’s medium with 1% non-essential amino acids; NEAA, Gibco). All cell culture media were supplemented with 10% fetal calf serum. Cells were passaged weekly and detached using trypsin. For imaging experiments, HEK STIM1/2^-/-^ were transfected via electroporation using the Amaxa Nucleofector II (Lonza) according to the manufacturer’s protocol. For co-immuniprecipitation, HEK STIM1/2^-/-^ cells were transfected using JetOptimus according to the manfucaturer’s protocol. SH-SY5Y cells were transfected using JetPrime (Polyplus transfections) transfection reagent according to the manufacturer’s protocol. Cells were analyzed 24h post transfection.

| Construct | Fluorescent marker | Vector |
| --- | --- | --- |
| HA-STIM2.2 | - | pEX-SP1 |
| HA-STIM2.3 | - | pEX-SP1 |
| mKate2-STIM2.2 | mKate2, N-term | pEX-SP1 |
| mKate2-STIM2.3 | mKate2, N-term | pEX-SP1 |
| mCherry-STIM1 | mCherry, N-term | pEX-SP1 |
| STIM2.2-mCherry | mCherry, C-term | pEX-SP1 |
| YFP-STIM2.2 | YFP, N-term | pEX-SP1 |
| YFP-STIM2.3 | YFP, N-term | pEX-SP1 |
| YFP-STIM2.2Δ5K | YFP, N-term | pEX-SP1 |
| YFP-STIM2.2Δ674 | YFP, N-term | pEX-SP1 |
| YFP-STIM2.2 2xIP | YFP, N-term | pEX-SP1 |
| YFP-STIM2.2 2xIP+Δ5K | YFP, N-term | pEX-SP1 |
| YFP-STIM2.2Δ711 | YFP, N-term | pEX-SP1 |
| YFP-STIM2.2Δ675-710 | YFP, N-term | pEX-SP1 |
| NFAT1-GFP | GFP | pCAGGS |
| HA-STIM2.2-YFPc | HA, N-term; YFPc, C-term | pEX-SP1 |
| STIM2.2-YFPn | YFPn, C-term | pBabe |
| HA-STIM2.3-YFPc | HA, N-term; YFPc, C-term | pEX-SP1 |
| STIM2.3-YFPn | YFPn, C-term | pBabe |
| STIM1-YFPn | YFPn, C-term | pBabe |
| Orai1-YFPn | YFPn, C-term | pMax |
| Orai2-YFPn | YFPn, C-term | pMax |
| Orai3-YFPn | YFPn, C-term | pMax |
| EFEMP1-YFPn | YFPn, C-term | pBabe |

### Generation of stable cell line

To generate a cell line stably expressing HA-STIM2.3, SH-SY5Y STIM2^-/-^ cells were detached using trypsin and seeded into 60 mm dishes containing DMEM + 10 % FCS + 1 % NEAA 24 hours prior to transfection using JetPrime according to the manufacturer’s protocol. 24 hours post transfection, medium was changed to DMEM + 10 % FCS + 1 % NEAA + 1mg/ml G418 as the selection antibiotic. The selection medium was changed regularly until constant growing was observed. The sensitivity of SH-SY5Y STIM2^-/-^ cells to G418 was determined beforehand using different concentrations ranging from 0,1 – 1mg/ml G418 on untransfected cells. The stable overexpression of HA-STIM2.2 or HA-STIM2.3 was analyzed using Ca^2+^ imaging and Western Blot.

### PCR and quantitative real-time PCR

RNA isolation was performed using TRIzol according to the manufacturer’s protocol. For cDNA synthesis, SuperScriptTM II Reverse Transcriptase (Life Technologies) was used. qRT-PCRs were performed using QuantiTect SYBR Green Kit (Qiagen) and a CFX96 Real-Time System (Bio-Rad). Primers are listet in. Results were normalized to TBP using the ΔC_q_ method. Values are shown as 2^-(ΔCq)^.

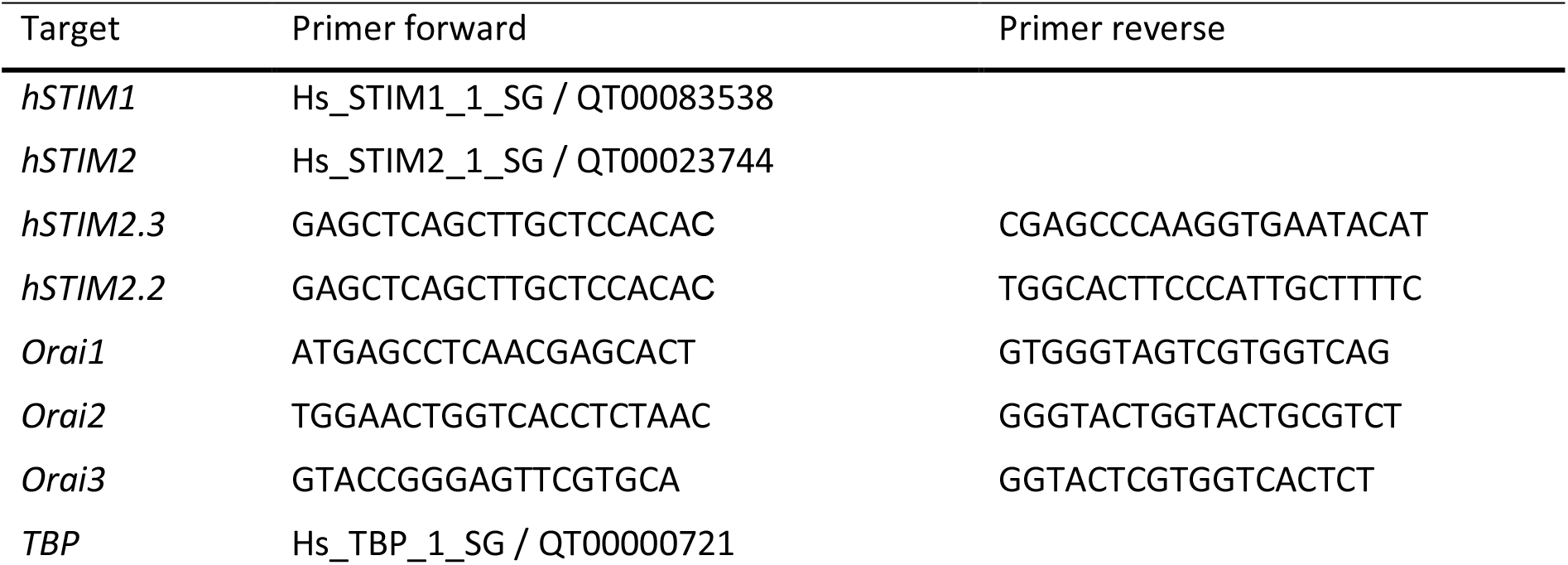

### RNA Sequencing

For RNA sequencing, RNA was isolated using TRIzol according to the manufacturer’s protocol and sent to Novogene (Cambridge, UK), who performed all further steps including the bioinformatics analysis. Differential gene expression analysis was performed using the DESeq2 R package for the anaylsis of two conditions or groups [52]. Subsequent Gene Ontology (GO) analysis using the clusterProfiler software [53] identifies associated cellular components, molecular functions and biological processes, which are significantly affected by the DEGs.

### Biochemistry

Transfected cells were washed with ice-cold PBS, detached in ice-cold RIPA buffer containing 150mM NaCl, 50mM Tris-HCl, 1% Non-Idet P40, 1% Triton and Protease Inhibitor Complete (Sigma-Aldrich) using a cell scraper. Cells were lysed for 5 min at -80°C, vortexed and centrifuged at 21380 g (4°C) for 20 min. For co-immuniprecipitation,transfected cells were washed in ice-cold PBS, detached in IP buffer containing 20mM Tris-HCl, 100mM KCl, 10% glycerol (v/v), 0,5% n-Dodecyl ß-D-maltoside (DDM) using a cell scraper. Cells were lysed for 30min at 4°C end-over-end tumbling and centrifuged at 18,000 g (4°C) for 30min. Protein concentrations were determined using BCA reagent according to the manufacturer’s protocol. Co-immunoprecipitation was performed overnight end-over-end tumbling at 4°C using protein low binding tubes and Pierce anti-HA agarose. The agarose was washed 4 times for 5 min in ice cold lysis buffer and centrifuged for 30 s at 500 g before being eluted with 50mM Tris-HCl pH 6.8 + 2%SDS for 5min at 90°C. Protein fractions were denatured in Laemmli buffer at 65°C for 15 min. Discontinous SDS-PAGE was performed with subsequent electrotransfer to nitrocellulose or PVDF membrane for 90 min at 150mV and 4°C. Membranes were blocked in 5% skimmed milk in TBS-T buffer for 60 min. Primary antibodies (α-HA3F10 Roche were incubated overnight at 4°C in PBS containing 1% BSA and 0.02% sodium azide. Membranes were washed 3 times for 10 min in TBS-T and incubated with the secondary antibody in blocking buffer for 60 min at RT. After 3 times washing for 10 min, the membranes were incubated in Clarity Western ECL Substrate and detection of chemiluminescence was performed on a CCD camera equipped ChemiDoc XRC+ system. Densitometric quantification of non-saturated signals was done using ImageLab software. Primary and secondary antibodies are listed in.

| Antigen | Manufacturer | Catalog # | clonality | Dilution |
| --- | --- | --- | --- | --- |
| α-phospho-AMPK | Cell signaling | 2535 | polyclonal | 1:1000 |
| α-AMPK | Cell signaling | 2532S | polyclonal | 1:1000 |
| α-β-Actin | Sigma Aldrich | A5441 | monoclonal | 1:5000 |
| α-HA 3F10 | Roche | 11867423001 | | 1:1000 |
| α-STIM1 | Proteintech | 11565-1-AP | polyclonal | 1:1000 |
| α-STIM2 | Alomone | ACC-064 | polyclonal | 1:300 |

| Antigen | Manufacturer | Catalog# | Dilution |
| --- | --- | --- | --- |
| α-rabbit-HRP | GE Healthcare/Amersham | NA9340 | 1:10000 |
| α-rat-HRP | Sigma Aldrich | A5795 | 1:20000 |
| α-mouse-HRP | Amersham/Bioscience | NA931 | 1:5000 |

### Fluorescence-based Ca imaging

HEK STIM1/2^-/-^ cells were seeded onto 25mm coverslips in 35mm dishes containing cell culture medium after transfection with 1μg YFP-/HA-/mKate2-tagged STIM2.2 or STIM2.3 (all pEX) using electroporation. SH-SY5Y STIM2^-/-^ or STIM1/2^-/-^ cells were seeded in 35mm dishes 24 hours prior to transfection with 2μg of YFP/HA-/mKate2-tagged STIM2.2 or STIM2.3 (all pEX). Transfected cells were reseeded 5 hours post transfection onto 25mm coverslips. After 16-24h, cells were loaded with 1μM Fura-2 in DMEM+10%FCS (SH-SY5Y: 1% NEAA) at RT for 30 min, transferred to a perfusion chamber and washed with external Ca^2+^ Ringer solution containing (in mM): 155 NaCl, 2 MgCl_2_, 10 glucose, 5 HEPES and 0.5-1.5 CaCl_2_ or no CaCl_2_, but 1 EGTA and 3 MgCl_2_ instead. ER stores were depleted using 1μM Thapsigargin (Tg) in 0 mM Ca^2+^ (Ca^2+^ readdition) or 0.5-1.5 mM Ca^2+^ (global) ringer solution. The imaging setup consisted of a Zeiss Axio Observer A1 inverse microscope, Polychrome V (Till Photonoics) light source and Clara CCD camera (Andor). F76-521 Fura-2 HC, F36-528 YFP and F20-451 RFP filter cubes (AHF Analysetechnik AG) were used. 340nm and 380nm images were acquired with 10-30ms exposure times and 1×1 or 2×2 pixel binning through a 20×0.75 Fluar (Zeiss) objective. Acquisition was controlled by and background-subtracted 340nm/380nm ratio values were calculated using VisiView Software. Ratio data was further analyzed using IgorPro (Wavemetrics). For kinetic parameters, average (5-10 frames) basal and plateau ratio values and maximal Tg-or Ca^2+^ readdition-induced peak ratio values were determined. To obtain Δs, the average ratio before addition of Tg or Ca^2+^ was subtracted, respectively. The influx rate is represented by the slope of a linear fit performed on the ratio values of 1 frame before and 4 frames after Ca^2+^ readdition. For quantification of global Ca^2+^ entry, the area under the curve upon store depletion was determined. The data was then imported into GraphPad 8.4 (Prism) for further statistical analysis and plotting.

### Live cell imaging

HEK STIM1/2^-/-^ were transfected with STIM2.2-mCherry and YFP-STIM2.3 or YFP-STIM2.2Δ5K or YFP-STIM2.2Δ674. Fluorescence was detected using the wide-field epifluorescence microscope cell observer A1 (Zeiss) with the Fluor 100x/ oil objective. To detect YFP, fluorescence filter cube 54HE and LED 470 Ex (470/40) were used, and for mCherry detection, filter cube 56HE and LED N-White + Ex (556/20) were used. Images were taken as Z-stacks and processed as a maximum intensity projection (MIP) for visualization. Colocalization was analyzed using Fiji plugin JACoP from a single stack. Threshold was set as mean plus two times SD.

### TIRF microscopy for cluster analysis

Cluster formation was monitored using a Leica TIRF MC system. HEKSTIM1/2 ^-/-^ cells were seeded into 35mm dishes 24 hours prior to transfection with PH-PLCγ-mCherry and YFP-STIM2.2 or YFP-STIM2.3 or YFP-STIM2.2Δ5K or YFP-STIM2.2 2xIP mutant+Δ5K. After 4 hours of incubation cells were reseeded onto 25mm coverslips in 35mm dishes containing DMEM +10% FCS. 24 hours after transfection, coverslips were transferred to a perfusion chamber and washed with 0.5 mM Ca^2+^ ringer solution. Store depletion and subsequent cluster formation was induced after 60s baseline acquisition by perfusion with 0.5 mM Ca^2+^ ringer solution containing 1 μM Thapsigargin. Images were taken with a 100×1.47 oil HCX PlanApo objective. YFP and mCherry were excited using 488nm or 561nm lasers with BP525/50 and DRT emission filters, respectively. The TIRF focal plane was set according to the PH-PLCγ-mCherry signal and a penetration depth of 90 nm was choosen.

### NFAT Assay

SH-SY5Y STIM1/2^-/-^ cells were seeded in 35mm dishes 24 hours prior to transfection with NFAT1-GFP and mKate2-STIM2.2, mKate2-STIM2.3, mKate2-STIM2.2Δ5K or mKate2-STIM2.2 2xIP+Δ5K. After 5 hours of incubation, cells were reseeded onto 25 mm coverslips in 35 mm dishes. 24 hours post transfection, NFTA1-GFP signal in the cytoplasm vs nucleus was detected on the Zeiss Axio Observer A7 equipped with a Prime95B sCMOS camera (Photometrics) using a 63x/1.40 Plan Apochromat oil objective. YFP and mKate2 signals were excited using an HXP 120 V compact light source with 480-498nm/560-584nm excitation filters and Fura8 (BS488nM, BP500-550nm)/63 HE Red (BP559-585nm, BS590nm, BP600-690nm) emission filter cubes. Cells were imaged in a Z-plane near the nucleus stained with 1mg/ml Hoechst 33342 dye immediately before the measurement. Cells were activated using 1 μM Thapsigargin in 1 mM Ca^2+^ ringer solution. The ratio of GFP signal in the cytoplasm vs. in the nucleus was determined using Fiji.

### BiFC

HEK STIM1/STIM2^-/-^_cells (Zhou et al, 2018) were transfected in a 6-well scale with HA-STIM2.2 or HA-STIM2.3 fused to aa 156–720 of YFP (YFPc) in pEX as bait in combination with POI fused to aa 1–155 of YFP (YFPn) in pBabe or pMax as prey. Transfection was performed at equimolar ratios of DNA using a total amount of 2 μg DNA. To induce SOCE, 1 μM TG was added to the media and incubated for 10 min 24 h post transfection. After detaching cells in PBS containing 1 μM TG, cells were centrifuged at 320 g in FACS tubes at 4°C for 10 min and resuspended in 400 μl ice-cold FACS buffer (5% FCS, 0.5% BSA and 0.07% NaN_3_ in PBS) with additional Zombie Aqua™ (1:1,000 Fixable Viability Kit, BioLegend) to stain for vital cells. FACSVerse system (BD Biosciences) was used to screen for YFP signal in vital cells. BD FACSuite™ and FloJo 10.0.7 (BD Biosciences) were used for determination of the percentages of YFP-positive cells.

### Bioinformatics

The following three datasets were downloaded from the NCBI Sequence Read Archives (SRA): SRP331938, GSE181813 representing 6 male and 6 female non-alcohol use disorder postmortem control donors, SRP346150: analyzing GSE207713 with iPSC derived astrocytes from 4 control donors, and GSE188847, which analyzes aged Covid-19 unaffected postmortem controls (, including 6 males, and 4 females), ages of age from 45 to 64 yrs., data years old. Additional datasets were obtained for development stages Carnegie stage 22 and 9PCW9 post conception weeks, downloaded from the Biostudies database, a data infrastructure belonging to The European Bioinformatics Institute https://www.ebi.ac.uk/biostudies/arrayexpress/studies/E-MTAB-4840 [54].

A bioinformatics workflow was constructed to first quantify gene expression and to analyze splicing events for selected genes. Files in FASTQ format for all datasets were downloaded and used as input for the nf-core *rnaseq* pipeline to map and quantify reads abundance using Salmon’s fast mapping algorithm [55]. Mapped reads in BAM format were sequentially subjected to splicing variations detection by MAJIQ software. Among the current tools for splice events study, MAJIQ is a comparably fast and reliable tool [56]. MAJIQ yields a Percent Selected Index (PSI) for each recognized and de-novo junctions as quantification measurements for splice variations, which efficiently facilitates our current study to identify and compare STIM2 alternative events across various samples. For both mapping and splicing analysis stages, the Human Reference Genome build 38 (hg38) retrieved from NCBI in GTF format was used. For easier comparison to qRTPCR data, single exon 13 splice border probabilities were averaged to give the overall exon 13 inclusion probability.

### Statistical analysis

Statistical analysis was performed using the software GraphPad Prism 9. The statistical test was chosen according to the normal distribution and number of conditions. Significances are as follows: * p<0.05, ** p<0.01, *** p<0.001. Data is displayed as mean ± SEM obtained from 3 independent transfections.

### Ethics votum

Ethical permission to obtain tissues from postmortem body donors was granted by the ethics commission of the medical board of the Saarland, Nrs. 163/20 and 148/18.

## Supporting information

Poth et al supplements

## Acknowledgements

We thank the Drs. Marcus Grimm and Tobias Hartmann, Saarland University for initial transfection of SH-SY5Y cells with the STIM2 targeting construct, Dr. Nicole Ludwig, Saarland University, for additional samples of RNA. Funding was provided by the Deutsche Forschungsgemeinschaft (DFG) grants SFB894 (A2) and SFB1027 (C4) to BAN.

## Author contribution

VP conducted all imaging and microscopy experiments, biochemistry and cell line generation, analyzed data and contributed to writing. HTTD and VH provided bioinfomatic analysis, KF performed RTPCR analysis, TT operated and provided postmortem tissues, DA helped with microscopy and initial experiments, BAN conceptualized the study, analyzed data and wrote the MS.

## References

1. Pan, Q., et al., Deep surveying of alternative splicing complexity in the human transcriptome by high-throughput sequencing. Nat Genet, 2008. 40(12): p. 1413–5.

2. Wang, E.T., et al., Alternative isoform regulation in human tissue transcriptomes. Nature, 2008. 456(7221): p. 470–6.

3. Ule, J. and B.J. Blencowe, Alternative Splicing Regulatory Networks: Functions, Mechanisms, and Evolution. Mol Cell, 2019. 76(2): p. 329–345.

4. Mazille, M., et al., Stimulus-specific remodeling of the neuronal transcriptome through nuclear intron-retaining transcripts. EMBO J, 2022. 41(21): p. e110192.

5. Siller, A., et al., beta2-subunit alternative splicing stabilizes Cav2.3 Ca(2+) channel activity during continuous midbrain dopamine neuron-like activity. Elife, 2022. 11.

6. Marasco, L.E. and A.R. Kornblihtt, The physiology of alternative splicing. Nat Rev Mol Cell Biol, 2022.

7. Gracheva, E.O., et al., Ganglion-specific splicing of TRPV1 underlies infrared sensation in vampire bats. Nature, 2011. 476(7358): p. 88–91.

8. Bell, L.R., et al., Positive autoregulation of sex-lethal by alternative splicing maintains the female determined state in Drosophila. Cell, 1991. 65(2): p. 229–39.

9. Brandman, O., et al., STIM2 is a feedback regulator that stabilizes basal cytosolic and endoplasmic reticulum Ca2+ levels. Cell, 2007. 131(7): p. 1327–39.

10. Stathopulos, P.B., L. Zheng, and M. Ikura, Stromal interaction molecule (STIM) 1 and STIM2 calcium sensing regions exhibit distinct unfolding and oligomerization kinetics. J Biol Chem, 2009. 284(2): p. 728–32.

11. Prakriya, M. and R.S. Lewis, Store-operated calcium channels. Physiological Reviews, 2015. 95: p. 1383–1436.

12. Williams, R.T., et al., Identification and characterization of the STIM (stromal interaction molecule) gene family: coding for a novel class of transmembrane proteins. Biochem J, 2001. 357(Pt 3): p. 673–85.

13. Niemeyer, B.A., Changing calcium: CRAC channel (STIM and Orai) expression, splicing, and posttranslational modifiers. Am J Physiol Cell Physiol, 2016. 310(9): p. C701–9.

14. Darbellay, B., et al., STIM1L is a new actin-binding splice variant involved in fast repetitive Ca 2+ release. Journal of Cell Biology, 2011. 194: p. 335–346.

15. Miederer, A.M., et al., A STIM2 splice variant negatively regulates store-operated calcium entry. Nature Communications, 2015. 6.

16. Rana, A., et al., Alternative splicing converts STIM2 from an activator to an inhibitor of store-operated calcium channels. Journal of Cell Biology, 2015. 209: p. 653–670.

17. Ramesh, G., et al., A short isoform of STIM1 confers frequency-dependent synaptic enhancement. Cell Rep, 2021. 34(11): p. 108844.

18. Knapp, M.L., et al., A longer isoform of Stim1 is a negative SOCE regulator but increases cAMP-modulated NFAT signaling. EMBO Rep, 2022. 23(3): p. e53135.

19. Xie, J., et al., Identification of a STIM1 Splicing Variant that Promotes Glioblastoma Growth. Adv Sci (Weinh), 2022. 9(11): p. e2103940.

20. Berna-Erro, A., et al., STIM2 regulates capacitive Ca2+ entry in neurons and plays a key role in hypoxic neuronal cell death. Sci Signal, 2009. 2(93): p. ra67.

21. Popugaeva, E., et al., STIM2 protects hippocampal mushroom spines from amyloid synaptotoxicity. Mol Neurodegener, 2015. 10: p. 37.

22. Sun, S., et al., Reduced synaptic STIM2 expression and impaired store-operated calcium entry cause destabilization of mature spines in mutant presenilin mice. Neuron, 2014. 82(1): p. 79–93.

23. Chanaday, N.L., et al., Presynaptic store-operated Ca(2+) entry drives excitatory spontaneous neurotransmission and augments endoplasmic reticulum stress. Neuron, 2021. 109(8): p. 1314–1332 e5.

24. Zhou, Y., et al., Cross-linking of Orai1 channels by STIM proteins. Proceedings of the National Academy of Sciences of the United States of America, 2018. 115: p. E3398–E3407.

25. Ong, H.L., et al., STIM2 enhances receptor-stimulated Ca(2)(+) signaling by promoting recruitment of STIM1 to the endoplasmic reticulum-plasma membrane junctions. Sci Signal, 2015. 8(359): p. ra3.

26. Pchitskaya, E., et al., Stim2-Eb3 Association and Morphology of Dendritic Spines in Hippocampal Neurons. Sci Rep, 2017. 7(1): p. 17625.

27. Ahmad, M., et al., Functional communication between IP(3)R and STIM2 at subthreshold stimuli is a critical checkpoint for initiation of SOCE. Proc Natl Acad Sci U S A, 2022. 119(3).

28. Gulyas, G., et al., LIPID transfer proteins regulate store-operated calcium entry via control of plasma membrane phosphoinositides. Cell Calcium, 2022. 106: p. 102631.

29. Subedi, K.P., et al., STIM2 Induces Activated Conformation of STIM1 to Control Orai1 Function in ER-PM Junctions. Cell Rep, 2018. 23(2): p. 522–534.

30. Chauhan, A.S., et al., STIM2 interacts with AMPK and regulates calcium-induced AMPK activation. FASEB J, 2019. 33(2): p. 2957–2970.

31. Ramamurthy, S., et al., AMPK activation regulates neuronal structure in developing hippocampal neurons. Neuroscience, 2014. 259: p. 13–24.

32. Williams, T., et al., AMP-activated protein kinase (AMPK) activity is not required for neuronal development but regulates axogenesis during metabolic stress. Proc Natl Acad Sci U S A, 2011. 108(14): p. 5849–54.

33. Florio, M., et al., Human-specific gene ARHGAP11B promotes basal progenitor amplification and neocortex expansion. Science, 2015. 347(6229): p. 1465–70.

34. Florio, M., et al., A single splice site mutation in human-specific ARHGAP11B causes basal progenitor amplification. Sci Adv, 2016. 2(12): p. e1601941.

35. Xing, L., et al., Expression of human-specific ARHGAP11B in mice leads to neocortex expansion and increased memory flexibility. EMBO J, 2021. 40(13): p. e107093.

36. Barton, R.A. and C. Venditti, Rapid evolution of the cerebellum in humans and other great apes. Curr Biol, 2014. 24(20): p. 2440–4.

37. Elorza, A., et al., Huntington’s disease-specific mis-splicing unveils key effector genes and altered splicing factors. Brain, 2021. 144(7): p. 2009–2023.

38. Smith, M.A., J. Brandt, and R. Shadmehr, Motor disorder in Huntington’s disease begins as a dysfunction in error feedback control. Nature, 2000. 403(6769): p. 544–9.

39. Singh-Bains, M.K., et al., Cerebellar degeneration correlates with motor symptoms in Huntington disease. Ann Neurol, 2019. 85(3): p. 396–405.

40. Vigont, V.A., et al., STIM2 Mediates Excessive Store-Operated Calcium Entry in Patient-Specific iPSC-Derived Neurons Modeling a Juvenile Form of Huntington’s Disease. Front Cell Dev Biol, 2021. 9: p. 625231.

41. Chang, C.L., et al., EB1 binding restricts STIM1 translocation to ER-PM junctions and regulates store-operated Ca(2+) entry. J Cell Biol, 2018. 217(6): p. 2047–2058.

42. Kim, J.H., et al., The TAM-associated STIM1(I484R) mutation increases ORAI1 channel function due to a reduced STIM1 inactivation break and an absence of microtubule trapping. Cell Calcium, 2022. 105: p. 102615.

43. Palty, R., et al., SARAF inactivates the store operated calcium entry machinery to prevent excess calcium refilling. Cell, 2012. 149(2): p. 425–38.

44. Zomot, E., et al., Bidirectional regulation of calcium release-activated calcium (CRAC) channel by SARAF. J Cell Biol, 2021. 220(12).

45. Jha, A., et al., The STIM1 CTID domain determines access of SARAF to SOAR to regulate Orai1 channel function. J Cell Biol, 2013. 202(1): p. 71–9.

46. Ahmad, M., et al., Functional communication between IP3R and STIM2 at subthreshold stimuli is a critical checkpoint for initiation of SOCE. Proc Natl Acad Sci U S A, 2022. 119(3).

47. Yoshida, T. and M. Mishina, Distinct roles of calcineurin-nuclear factor of activated T-cells and protein kinase A-cAMP response element-binding protein signaling in presynaptic differentiation. J Neurosci, 2005. 25(12): p. 3067–79.

48. Kar, P., et al., The N terminus of Orai1 couples to the AKAP79 signaling complex to drive NFAT1 activation by local Ca(2+) entry. Proc Natl Acad Sci U S A, 2021. 118(19).

49. Stein, B.D., et al., Quantitative In Vivo Proteomics of Metformin Response in Liver Reveals AMPK-Dependent and -Independent Signaling Networks. Cell Rep, 2019. 29(10): p. 3331–3348 e7.

50. Nelson, M.E., et al., Phosphoproteomics reveals conserved exercise-stimulated signaling and AMPK regulation of store-operated calcium entry. EMBO J, 2019. 38(24): p. e102578.

51. Poth, V., M.L. Knapp, and B.A. Niemeyer, STIM proteins at the intersection of signaling pathways. Current Opinion in Physiology, 2020. 17: p. 63–73.

52. Anders, S. and W. Huber, Differential expression analysis for sequence count data. Genome Biol, 2010. 11(10): p. R106.

53. Yu, G., et al., clusterProfiler: an R package for comparing biological themes among gene clusters. OMICS, 2012. 16(5): p. 284–7.

54. Lindsay, S.J., et al., HDBR Expression: A Unique Resource for Global and Individual Gene Expression Studies during Early Human Brain Development. Front Neuroanat, 2016. 10: p. 86.

55. Patro, R., et al., Salmon provides fast and bias-aware quantification of transcript expression. Nat Methods, 2017. 14(4): p. 417–419.

56. Mehmood, A., et al., Systematic evaluation of differential splicing tools for RNA-seq studies. Brief Bioinform, 2020. 21(6): p. 2052–2065.

