## Supplementary material for "A better brain? Alternative spliced STIM2 in hominoids arises with synapse formation and creates a gain-of-function variant": Poth et al supplements

### Figure S1

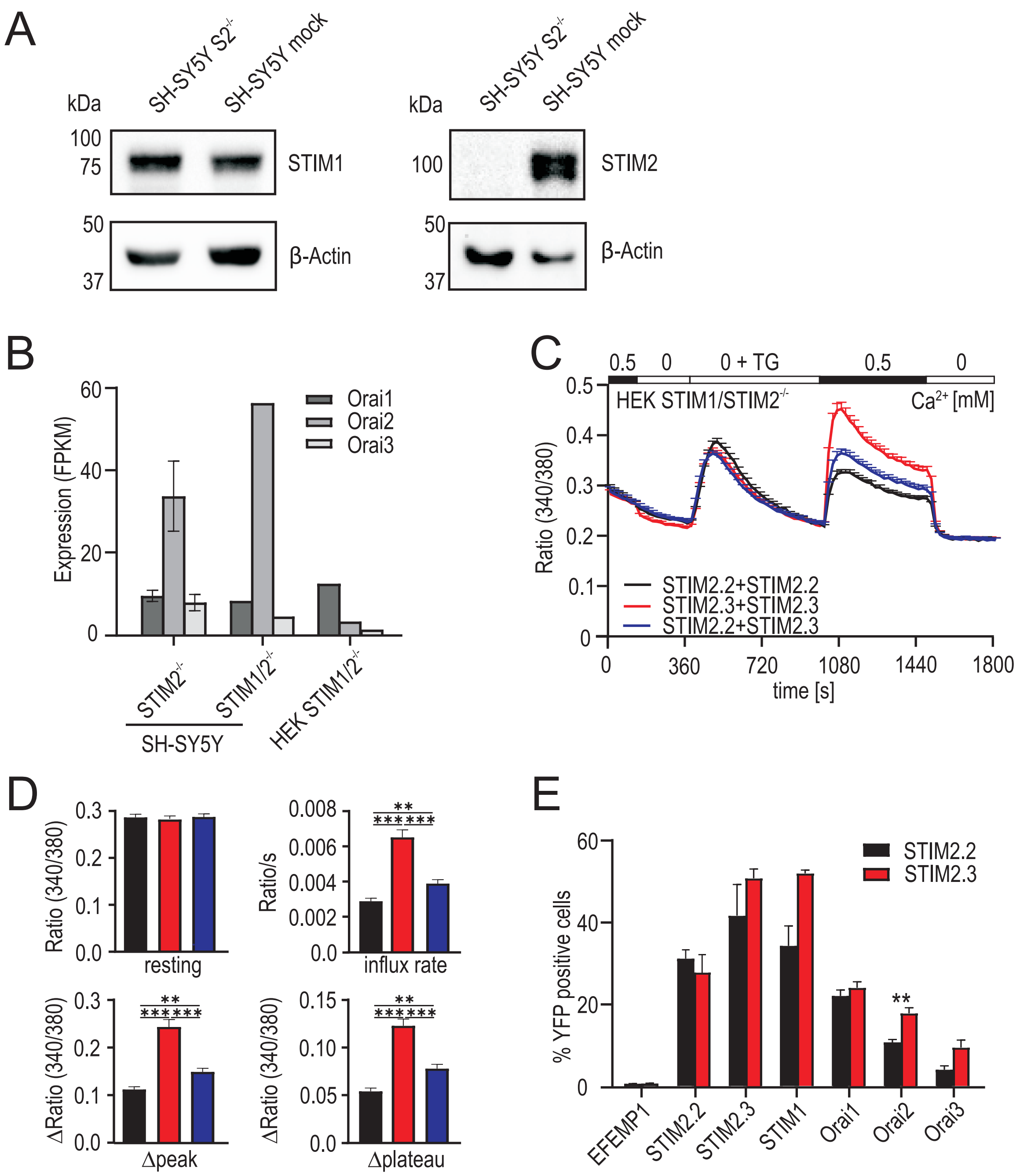


**Figure S1**

A. Western blot demonstrating absence of STIM2 in SH-SY5Y STIM2 $^{-/-}$.

B. FPKM values of *Orai 1-3* genes in SH-SY5Y STIM2 $^{-/-}$, SH-SY5Y STIM1/2 $^{-/-}$ and HEK STIM1/2 $^{-/-}$ cells from RNA Seq analysis.

C. Average traces showing changes (mean+SEM) in intracellular $Ca^{2+}$ (Ratio 340/380) over time in response to perfusion of different external $Ca^{2+}$ [mM] as indicated in the upper bar in HEK STIM1/2 $^{-/-}$ cells transfected with YFP-STIM2.2+mKate-STIM2.2 (black, n=100), YFP-STIM2.3+mKate-STIM2.3 (red, n=77) and YFP-STIM2.2+mKate-STIM2.3 (blue, n=99).

D. Quantification of changes in resting $Ca^{2+}$, influx rate, Δpeak and Δplateau measured in C. *** p< 0.001, ** p< 0.01; Kruskal-Wallis ANOVA.

E. Interaction of STIM2.2-YFPc (black) or STIM2.3-YFPc (red) with POI-YFPn in HEK STIM1/2 $^{-/-}$ cells was quantified as % YFP positive cells with bimolecular fluorescence complementation via flow cytometry. Data (mean+SEM) was obtained from 3 independent transfections with 10,000 measured cells each.

### Figure S2

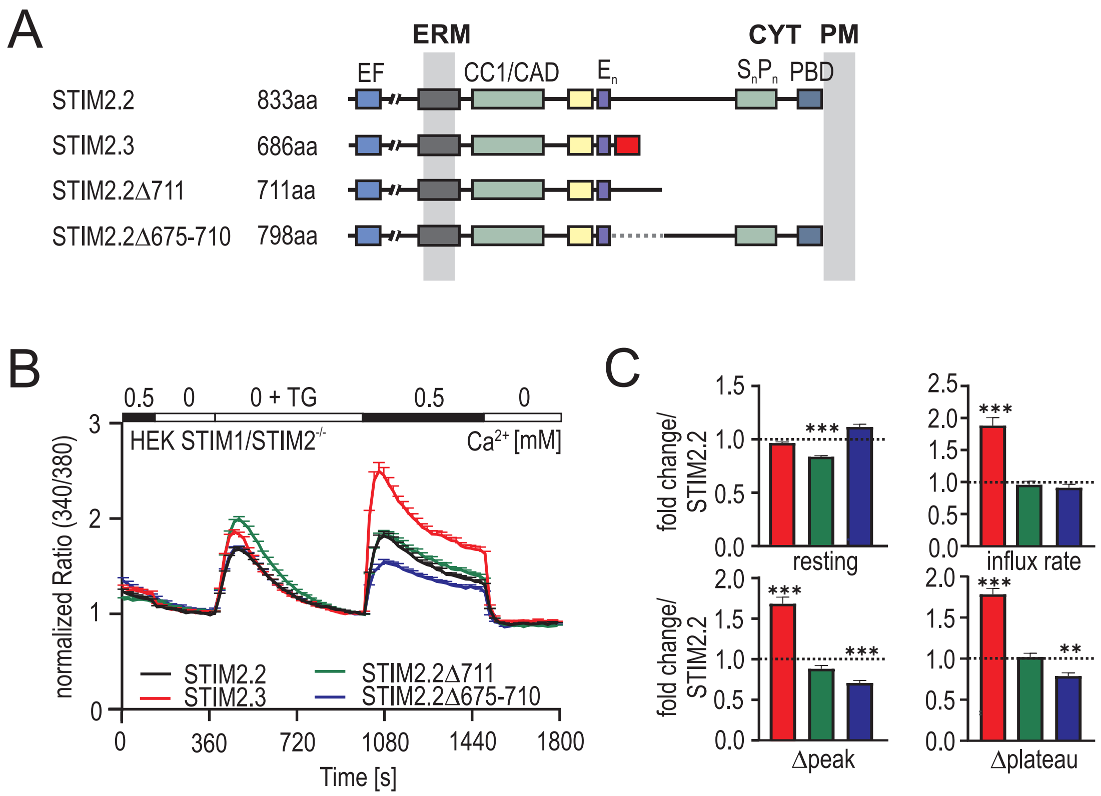


**Figure S2**

A. Schematic protein structure with functional domains of STIM2.2, STIM2.3, STIM2.2Δ711 terminating after aa 711 and STIM2.2Δ675 -710 with an internal deletion of 31 aa.

B. Normalized average traces of intracellular $Ca^{2+}$ (Ratio 340/380) over time in response to perfusion with different external $Ca^{2+}$ [mM] as indicated in the upper bar after transfection with YFP -STIM2.2 (black, n=165), YFP -STIM2.3 (red, n=207), YFP -STIM2.2 Δ711 (green, n=187) or YFP -STIM2.2Δ675 -710 (blue, n =117) in HEK STIM1/2 $^{-/-}$ cells.

C. Quantification of changes in resting $Ca^{2+}$, influx rate, Δpeak and Δplateau measured in B as fold change normalized to STIM2.2. *** $p< 0.001$, ** $p< 0.01$; Kruskal -Wallis ANOVA.

### Figure S3

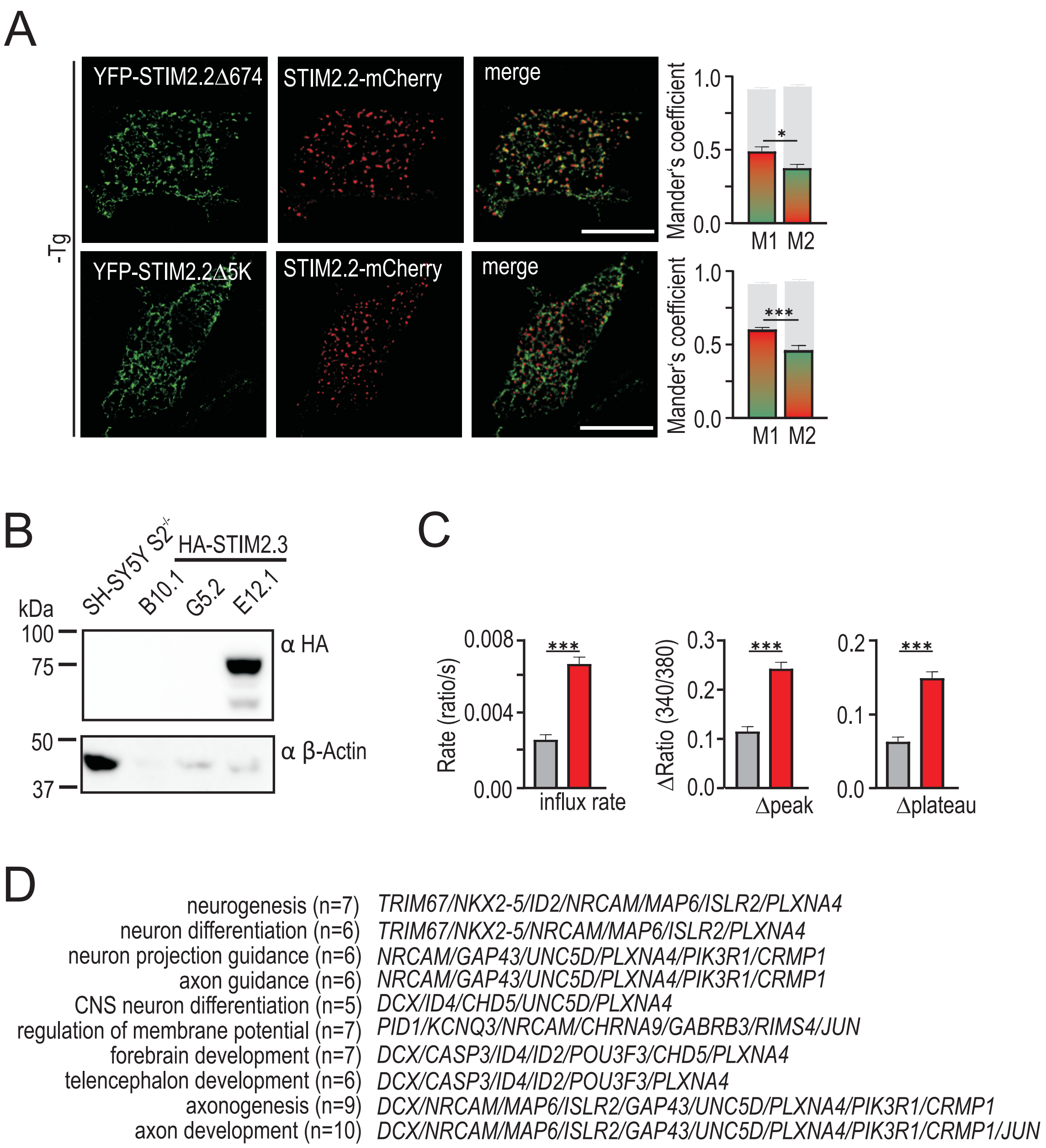


**Figure S3**

A. Representative images of HEK STIM1/2 $^{-/-}$ cells co-expressing STIM2. 2-mCherry (red) and YFP - STIM2.2 Δ674 (green) (upper panel) or STIM2.2 Δ5K (green) (lower panel) and merged images before Thapsigargin ( -Tg). Scale bar indic ates 10µM. For each condition 16 -29 cells from 3 independent transfections were analyzed using Mander's overlapping coefficients (M1, M2): STIM2.2+ Δ674 M1: 0.48 M2: 0.37; STIM2.2+ Δ5K M1: 0.58 M2: 0.45. Mander's overlapping coefficients of YFP -STIM2.2 and S TIM2.2-mCherry transfected cells are indicated by grey bars, M1: 0.90 M2: 0.92. * $p < 0.05$; *** $p < 0.001$, Mann -Whitney test.

B. Western Blot demonstrating over -expression of HA -STIM2.3 in the SH -SY5Y STIM2 -/- clone E12.1.

C. Quantification of influx rate, ✱peak and ✱plateau from cells measured in Fig. 7E. *** $p < 0.001$, Mann -Whitney test .

D. Corresponding genes of the identified GO terms in Fig. 7F.
